# Maternal-fetal immune conflict contributes to male-specific impairments in a mouse model of neurodevelopmental disorders

**DOI:** 10.64898/2026.02.02.703351

**Authors:** Irene Sanchez-Martin, Bharti Kukreja, Paige Henderson, Qianyu Lin, Daniel DiMartino, Valerie Bagan, Justin Park, Brian T. Kalish, Lucas Cheadle

## Abstract

Autism spectrum disorder (ASD) arises from genetic and environmental risk factors. One environmental factor, maternal immune activation (MIA)—in which pathogenic infection during pregnancy increases ASD risk in offspring—disproportionately affects males. However, the basis for this male-specific vulnerability, and the mechanisms by which inflammatory signals cross the maternal-fetal interface to affect the male embryo, remain largely unknown. Using the poly(I:C) mouse model of neurodevelopmental disorders, we characterize fetal, placental, and amniotic changes occurring within twenty-four hours of MIA. We find that 30% of embryos exhibit large-scale teratogenic abnormalities—ranging from decreased fetal weight to altered sensory organ development—while 70% develop normally. These abnormalities occur exclusively in a subset of males, never in females. Single-nucleus transcriptomics revealed robust induction of pro-inflammatory gene programs in the placentas of affected males across a broad range of cell types, and most prominently in spongiotrophoblasts—fetally derived cells that partly form the maternal-fetal border. These cells simultaneously downregulate extracellular matrix and hormone biosynthesis pathways, coinciding with a breakdown in placental structural integrity and the accumulation of immune cells and cytokines in the amniotic fluid. One such cytokine, IL-6, is required for MIA-evoked developmental abnormalities to emerge. Our data indicate that MIA drives a rapid shift from an immunosuppressive to a pro-inflammatory maternal-fetal interface in a vulnerable subset of male embryos, producing acute, sex-restricted developmental deficits. We propose that male embryos may express unique proteins capable of triggering this inflammatory response, which—combined with MIA-induced loss of maternal immunosuppression—selectively derails male embryonic development.

## INTRODUCTION

Autism spectrum disorder (ASD) is a heterogeneous class of neurodevelopmental conditions characterized by cognitive and social impairments that arises in early childhood and persists across the lifespan. ASD prevalence has increased substantially over recent decades, underscoring its growing impact on public health(*1–3*). A striking feature of ASD is that, while genetic factors play a strong role(*4*), environmental factors such as prenatal inflammation also contribute to neurodevelopmental dysfunction in a subset of individuals. For example, some epidemiological studies have identified an association between maternal exposure to certain pathogens, such as rubella, during pregnancy and heightened risk of neurodevelopmental disorders, such as ASD and schizophrenia, in the offspring(*5–8*). Despite this association, how a transient inflammatory insult during gestation compromises brain function across the lifespan remains poorly understood.

While prenatal inflammation is a major risk factor for ASD, the majority of fetuses exposed to inflammation in the womb undergo neurotypical development. This is consistent with resilience to maternal immune activation (MIA) existing across much of the population(*9*), with only a subset of individuals susceptible to acquiring ASD downstream of MIA. For example, even though epidemiological and animal studies suggest that ASD in females may present differently from its presentation in males(*10–15*), males remain more likely by about 4:1 to receive an ASD diagnosis(*16*). While males have been suggested to be more susceptible than females to MIA for reasons that remain a topic of debate(*12, 17*), one possibility is that male fetuses express paternally derived proteins that could cause the maternal immune system to tolerate the semi-allogeneic male fetus less well than the fetuses of females(*18, 19*). In turn, this loss of immune tolerance could disrupt neuro-immune homeostasis in the fetus. However, while this hypothesis is compelling, the factors that underlie the increased susceptibility of males to MIA remain to be established.

Mouse models of MIA have been instrumental in linking prenatal inflammation to ASD-relevant outcomes(*8*). A widely used paradigm involves administration of the viral mimetic poly(I:C) during midgestation. Prior work has focused primarily on behavioral consequences in adult offspring, unveiling ASD-relevant phenotypes including decreased sociability, an overabundance of repetitive behaviors, and heightened anxiety compared to saline-treated control mice(*20–23*). Moreover, several studies have reported cellular and molecular changes that occur at the maternal-fetal interface in response to poly(I:C), such as increased levels of the cytokines IL-6 and TNFα and the activation of macrophages and natural killer (NK) cells (*17, 24–27*). However, as most of these studies were performed several days after poly(I:C) injection, the earliest inflammatory events that trigger developmental impairments in the embryo downstream of MIA remain incompletely understood. Here, we harness the MIA_poly(I:C)_ model to shed light on the factors that underlie the susceptibility or resilience of individual embryos to MIA, uncovering evidence that maternal immune conflict with male embryos contributes to male-specific developmental deficits in this model of ASD. Our results lay the groundwork for studies that could inform the development of strategies for preventing or treating ASD in prenatal males.

## RESULTS

To uncover the mechanisms linking maternal inflammation to neurodevelopmental dysfunction, we adopted the MIA_poly(I:C)_ mouse model. In this model, pregnant mice are intraperitoneally administered 20 mg/kg of poly(I:C)(Sigma-Aldrich P9582, lots 2137379, 329448, and 373973), a synthetic double stranded RNA that predominantly (but not exclusively) binds Toll-like receptor 3 (TLR3) to induce the same innate immune pathways as a viral infection(*28, 29*). Administration of poly(I:C) (or saline as control) occurs at E12.5, a gestational timepoint that correlates with the late second trimester in humans, one of the gestational ages at which maternal infections have been linked to ASD in the offspring(*30, 31*). Yet, while the impact of prenatal inflammation on the adult offspring of MIA_poly(I:C)_ mice has been broadly assessed at the behavioral level, the initial mechanisms by which maternal inflammation triggers life-long deficits in brain function in the fetus remain largely unknown.

### MIA elicits large-scale developmental abnormalities in a subset of embryos within 24 hours of poly(I:C) exposure

To identify the earliest effects of MIA on fetal development, we administered poly(I:C) or saline to pregnant C57BL/6J dams at E12.5 then harvested the embryos 24 hours later (i.e. E13.5; Fig. 1A). Since MIA models exhibit substantial variability in their activation of the maternal immune system depending upon factors ranging from the mouse strain to the commercial source of poly(I:C)(*32–34*), we validated induction of inflammation in all dams used in the study by performing ELISAs to ensure that levels of IL-6 in the maternal blood are upregulated following poly(I:C) injection (Fig. 1B,C). The vast majority of dams exhibited the expected increase in IL-6; the ∼2% that did not were excluded from the study. Twenty-four hours later at E13.5, we harvested embryos, then applied a systematic battery of morphological measurements to assess differences between the groups, with all embryos from each litter included in the study for the purpose of comprehensive sampling. While embryos harvested from saline-treated mice closely resembled those of uninjected wild-type mice (Fig. S1A), embryos from poly(I:C)-treated dams displayed significant abnormalities. These abnormalities included changes in overall weight and morphology (Fig. 1D-I), craniofacial and head malformations such as decreased lambda-to-bregma distance (Fig. 1J-L), evidence of a lack of vascular integrity on the surface of the embryo (Fig. 1M), a lack of external sensory organ (i.e. ear pavilion and whisker pad) development (Fig. 1N), and abnormalities in spine and tail morphology (Fig. 1O). Although these differences between groups were statistically significant, the distributions of the data overlap substantially, suggesting that some fetuses were deleteriously impacted by MIA while others maintained a normal developmental trajectory. Thus, we next quantified the proportions of embryos per litter displaying at least one of these abnormalities between saline and poly(I:C) groups. Consistent with the partial overlap between the data distributions, we found that about 30% of embryos displayed one or more developmental abnormalities following MIA, versus 0% of embryos from the control group (Fig. 1P). Importantly, we observed the same effects when we applied the MIA_poly(I:C)_ paradigm to another well-studied but genetically distinct laboratory mouse strain, BALB/c mice, indicating that, while untested mouse strains may respond to MIA differently(*35*), our observations are not limited to mice of a single genetic background (Fig. S1B,C).

**Figure 1.**
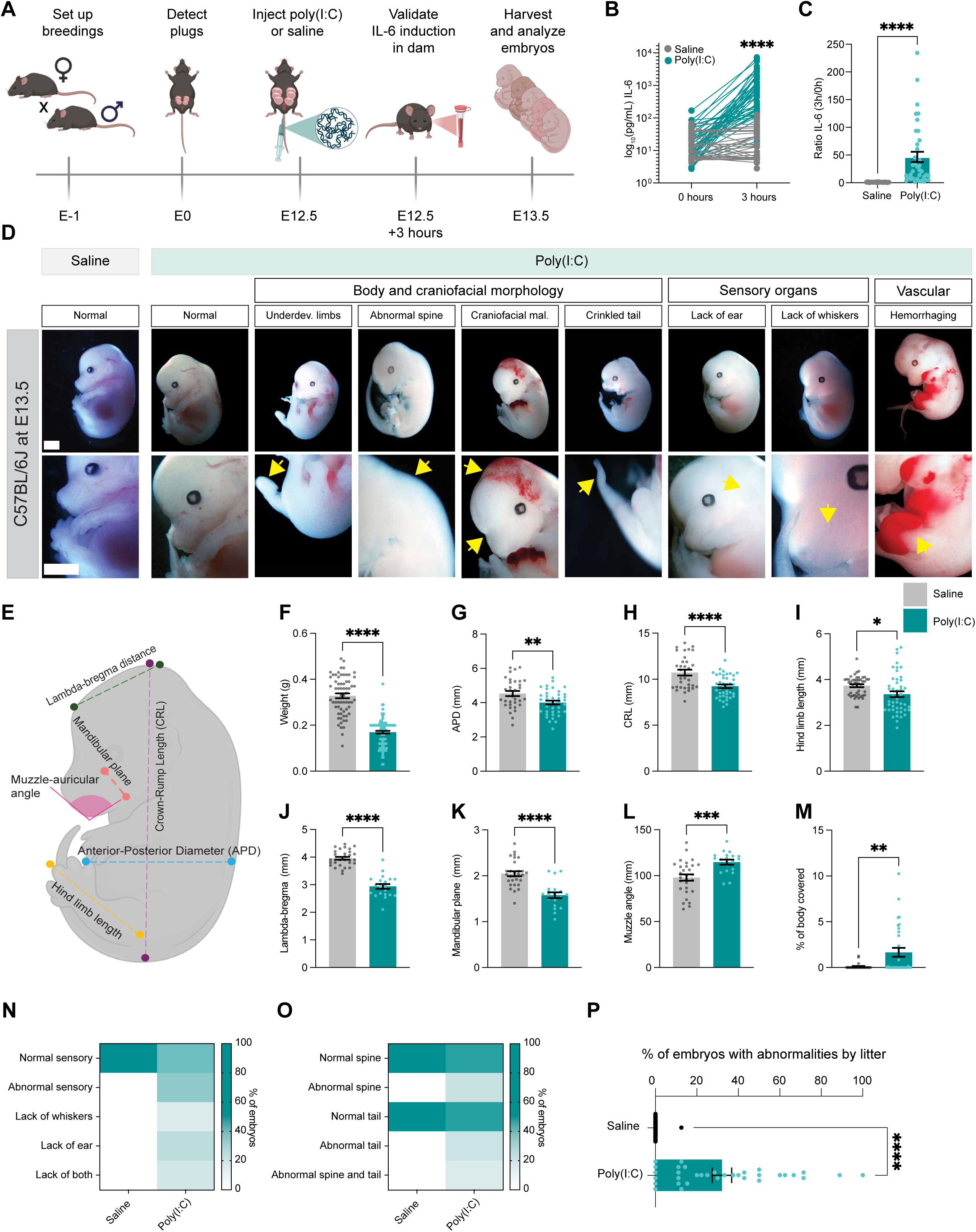
MIA elicits large-scale developmental abnormalities in a subset of embryos within 24 hours of poly(I:C) exposure. **(A)** Maternal immune activation paradigm. Created in BioRender. Cheadle, L. (2026) https://BioRender.com/iahyw0n. (**B**) IL-6 levels in the serum of dams exposed to poly(I:C) (teal) or saline (gray) measured by ELISA. Unpaired t test, ****p < 0.0001; n = 55 mice/condition. (**C**) Ratio of IL-6 levels before and three hours after poly(I:C) injection. Unpaired t test, ****p < 0.0001; n = 55 mice/condition. (**D**) Images of embryos harvested at E13.5. Yellow arrows indicate deficits observed. Underdev., underdeveloped; mal., malformation. Scale bars, 2 mm. (**E**) Schematic illustrating the nomenclature used to describe morphological dimensions of a fetus. Created in BioRender. Cheadle, L. (2026) https://BioRender.com/938kphb (**F**)-(**M**) Measurements following MIA: (**F**) fetal weight (g); (**G**) Anterior-posterior diameter (mm); (**H**) Crown-rump length (mm); (**I**) Hind limb length (mm); (**J**) Lambda-bregma distance (mm); (**K**) Mandibular plane length (mm); (**L**) Muzzle-auricular angle (mm); (**M**) Percent of body covered by hemorrhages. (**F**)-(**M**) Unpaired t tests, *p < 0.05; **p < 0.01; ***p < 0.001; ****p < 0.0001. For (**F**), n = 63 saline and 103 poly(I:C) embryos collected from 10 and 19 distinct litters, respectively. For (**G**)-(**H**), n = 33 saline and 40 poly(I:C). For (**I**), n = 28 saline and 40 poly(I:C). For (**J**)-(**L**), n = 30 saline and 21 poly(I:C). For (**M**), n = 35 embryos of each condition. (**N**) Heatmap displaying the percentage of embryos exhibiting normal or abnormal development of the ear pavilion and whisker pad. Scale on right. (**O**) Percentage of embryos exhibiting normal or abnormal development of the tail and spine. Scale on right. For (**N**),(**O**), n = 35 embryos acquired from 6 litters/condition. (**P**) Percentages of saline or poly(I:C)-exposed fetuses displaying one or more developmental abnormality by litter. Unpaired t test, ****p < 0.0001; n = 34 litters/condition.

### MIA-induced deficits in embryonic development are restricted to a subpopulation of male embryos

As individual embryos were differentially affected by MIA, we next classified the embryos of poly(I:C)-treated dams into three categories based upon the changes observed. Embryos were classified as those appearing indistinguishable from healthy controls (*normal*, poly(I:C)^norm^), those exhibiting at least one developmental abnormality but not showing signs of reabsorption (*moderate*, poly(I:C)^mod^), and those exhibiting extreme deficits which included signs of reabsorption, indicating that these embryos likely undergo intrauterine demise prior to birth (*strong*, poly(I:C)^strong^; Fig. 2A). To provide unbiased justification for this classification system, we performed a multiple correspondence analysis (MCA) using a panel of phenotypic descriptors encoded as binary indicators. This analysis shows a clear, graded separation of embryos into these three phenotypic groups along two dimensions. Dimension 1 captures the severity gradient such that poly(I:C)^strong^ cases cluster with traits like head and limb abnormality and absence of ears and whiskers. Poly(I:C)^norm^ embryos cluster at the opposite end, and poly(I:C)^mod^ embryos are distributed between these two groups. In contrast, dimension 2 captures additional phenotypic heterogeneity independent of overall severity. Specifically, embryos with intermediate versus severe phenotypes separated along this axis despite partial overlap in specific developmental abnormalities, suggesting that dimension 2 reflects differences in the combination and distribution of defects rather than a simple increase in severity. Together, these dimensions indicate that developmental abnormalities occur as coordinated multivariate patterns rather than isolated traits, supporting the biological relevance of the three-group classification system (Fig. 2B). These results are consistent with the observation that pathogenic infection in humans impacts fetal development heterogeneously, as well as the known link between prenatal inflammation and stillbirth.

**Figure 2.**
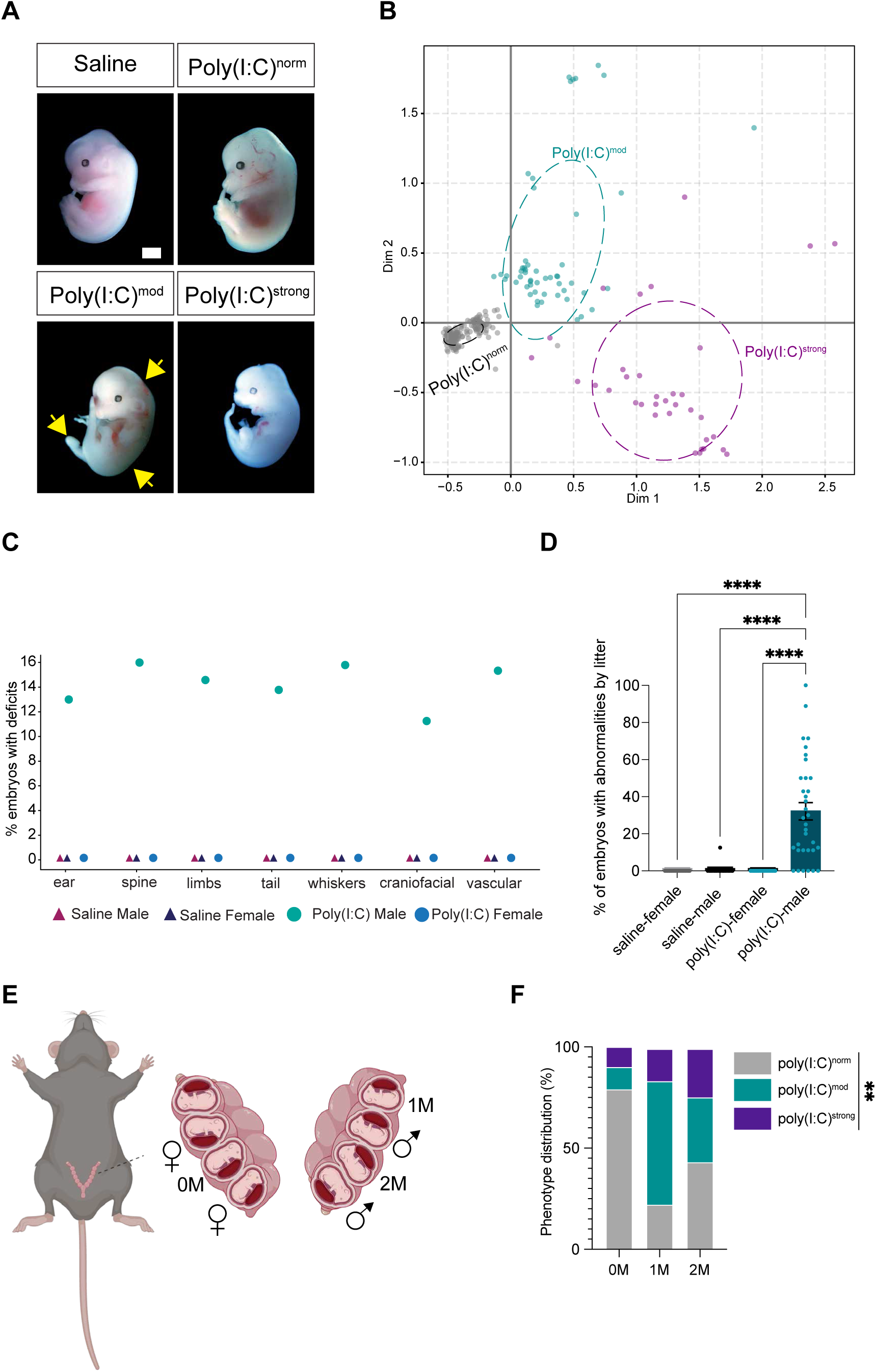
MIA-induced deficits in fetal development are restricted to a subset of male embryos. (**A**) Images of embryos labeled based upon phenotype following MIA: poly(I:C)^norm^, no abnormalities observed; poly(I:C)^mod^, at least one abnormality and no signs of reabsorption observed; and poly(I:C)^strong^, at least one abnormality and signs of reabsorption observed. Yellow arrows, underdeveloped limb and morphological abnormalities. Scale bar, 2 mm. (**B**) Multiple correspondence analysis (MCA) highlighting the separation of embryos into three distinct groups based upon measured phenotypes. Ellipses are centered at the group mean and oriented by the sample covariance. (**C**) Graph illustrating the percentage of embryos exhibiting deficits across both sexes and poly(I:C) or saline conditions. (**D**) Percentages of saline or poly(I:C)-exposed male and female fetuses displaying one or more developmental abnormalities plotted by litter. Two-Way ANOVA with Tukey’s post test, ****p < 0.0001. Treatment: *p < 0.05; Sex: ****p < 0.0001; Interaction: *p < 0.05. (**E**) Schematic illustrating the organization of embryos within the womb. Embryo surrounded only by female embryos (0M), embryo surrounded by one male embryo (1M), and embryo surrounded by two males (2M). Created in BioRender. Cheadle, L. (2026) https://BioRender.com/6pahnyu. (**F**) Quantification of phenotype distribution in 0M, 1M, and 2M embryos. Statistics; **p < 0.01.

We harnessed this classification system to identify factors that render some embryos susceptible to teratogenic deficits in response to MIA while others remain resilient. Since we inject poly(I:C) into the lower right quadrant of the dam’s abdomen, we first considered the possibility that embryos closest to the injection site are exposed to a greater concentration of inflammatory factors, thereby causing these embryos to undergo developmental changes more readily. However, we did not observe a significant correlation between phenotypic severity and the locus of injection (Fig. S2A,B).

We next considered the possibility that, even at this early developmental timepoint, the sex of the embryo might influence susceptibility to MIA. We used PCR to detect the Y chromosome genes *Rbm31* and *Sry* in poly(I:C)-exposed embryos across phenotypes (Fig. S2C). Strikingly, while the unaffected poly(I:C)^norm^ group included both male and female embryos, 100% of embryos within the poly(I:C)^mod^ and poly(I:C)^strong^ groups were male. Thus, roughly 60% of male fetuses from MIA dams exhibit some combination of developmental deficits at E13.5 compared to 0% of females (Fig. 2C). When plotted by litter, we find that on average about 30% of male embryos within a given litter exhibit deficits, although the distribution is wide, with some litters including no impacted embryos while others contain greater than 60% affected male embryos (Fig. 2D). The deleterious effect of MIA on male but not female embryonic development in C57Bl/6J mice occurs in BALB/c mice as well (Fig. S3). This finding is intriguing given the strong sex bias in disorders such as ASD, where males are more likely to receive a diagnosis than females at a ratio of 4:1(*16*). Thus, sex (but not intrauterine position) is one factor underlying risk of embryonic developmental dysfunction following MIA.

We next considered the possibility that the proximity of a male embryo to other embryos in the womb might influence its phenotype, as this relationship was recently shown to impact behavioral outcomes of MIA offspring(*36*). We found that embryos were significantly more likely to fall within the poly(I:C)^mod^ group (versus the poly(I:C)^norm^ group) if they were neighbored by one or two other male embryos than if they were surrounded by females. A similar pattern emerged for fetuses within the poly(I:C)^strong^ group (Fig. 2E,F). Thus, at least two factors underlie the selective susceptibility of a fetus to early developmental dysfunction in MIA: the sex of the fetus, and the sexes of neighboring fetuses in the womb.

### Inducing MIA after E12.5 fails to elicit developmental deficits

Prior work in mouse models of MIA suggests that the effects of poly(I:C) exposure on neurodevelopment can vary depending upon the stage of prenatal development at which inflammation is induced(*37*). Thus, we next asked whether E12.5 - E13.5 represents a period of development when some fetuses are particularly sensitive to inflammation. In support of this idea, when we injected poly(I:C) at E14.5 then assessed embryos at E15.5, fetal development was generally unaffected in both males and females (Fig. S4A-C). Conversely, inducing MIA at E9.5 led to developmental abnormalities or intrauterine demise in embryos of both sexes (Fig. S4D-F). These data suggest that the developmental age at which MIA occurs strongly influences its impact on fetal development, and that E12.5-E13.5, during which the placental barrier is still undergoing developmental changes in cellular composition and organization(*38, 39*), is a sensitive period of susceptibility of some male fetuses to MIA.

### The extracellular matrix is compromised in the placentas of poly(I:C)^mod^ embryos

C57Bl/6J mice typically carry litters of four to eight embryos at once, and each embryo harbors a unique maternal-fetal interface in the form of the placenta. The placenta is comprised of three layers: (1) the decidua which is derived from the uterus and is thus of maternal origin; (2) the fetally-derived spongiotrophoblast (SpT) layer which harbors the junctional zone, where maternal and fetal compartments of the placenta meet; and (3) the labyrinth, where maternal and fetal blood vessels intermingle and exchange nutrients, hormones, and other factors(*38, 40, 41*)(Fig. 3A). Given that poly(I:C) exposure induces developmental deficits in a subset of male embryos at E12.5, we hypothesized that MIA-induced changes in the placenta underlie the phenotypes we observe. To test this hypothesis, we performed bulk RNA-sequencing on the placentas of saline-treated male embryos and poly(I:C)-treated embryos that displayed developmental deficits without signs of reabsorption (i.e. the poly(I:C)^mod^ population; Fig. S5A). We find that genes encoding extracellular matrix (ECM) proteins, particularly those in the collagen (e.g. *ColGalT2*, *Col15a1, Col16a1, Col27a1, and Col12a1*) and laminin (*Lama3*, *Lamb3*, and *Lamc3*) families show reduced expression in poly(I:C)^mod^ embryos compared to controls (Fig. S5B,C). Our data suggest that the ECM is compromised in the placentas of male embryos that exhibit deficits in response to MIA.

**Figure 3.**
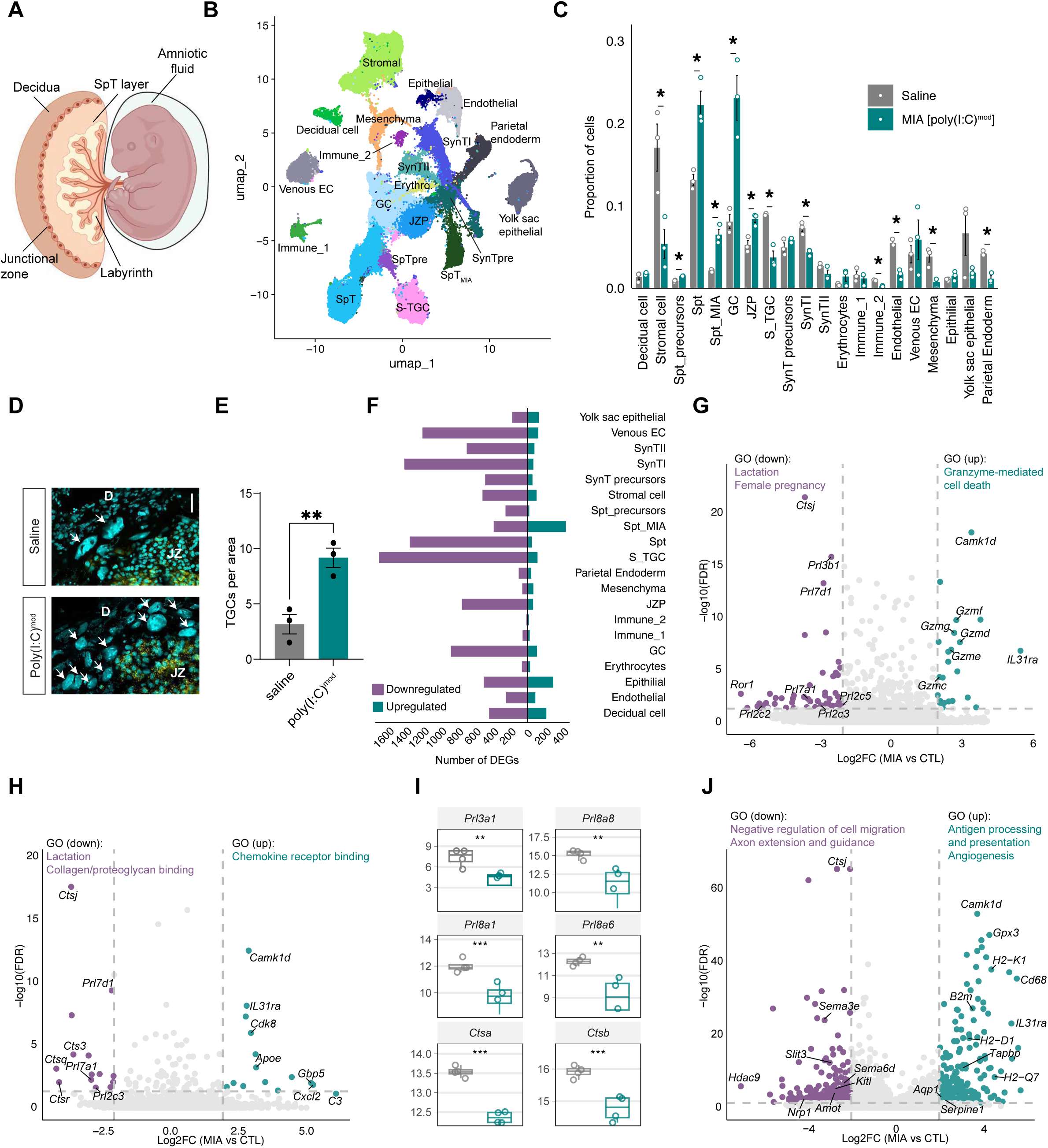
Single-nucleus RNA-sequencing unveils maternal and fetal immune activation in poly(I:C)^mod^ placentas. (**A**) Schematic of the tri-layered placenta with amniotic sac, amniotic fluid, and fetus shown. Created in BioRender. Cheadle, L. (2026) https://BioRender.com/0sfnngj (**B**) Uniform manifold approximation and projection (UMAP) plot illustrating the 197,237 nuclei included in the single-nucleus RNA-sequencing (snRNAseq) dataset, clustered by cell type. (**C**) Bar graph comparing the proportion of cells belonging to each cluster in saline-treated male (gray) and poly(I:C)^mod^ (teal) placentas. Independent two-sample t tests, *p < 0.05. (**D**) Confocal images of placental sections subjected to fluorescence *in situ* hybridization for the SpT marker *Ascl2* (yellow) with DAPI shown in cyan. D, decidua; JZ, junctional zone; arrows, TGCs. Scale bar, 50 µm. (**E**) Quantification of the number of TGCs per unit area in both conditions. Unpaired t test, **p < 0.01, n=3 placentas per condition. (**F**) The abundance of differentially expressed genes (DEGs) between conditions for each cell type meeting a threshold of 4.0 absolute fold change and FDR < 0.05. (**G**) Volcano plot illustrating DEGs identified in maternally derived placental immune cells. (**H**) Volcano plot illustrating DEGs identified in fetally derived immune cells. (**I**) RNA levels of each given gene as measured by bulk RNA-sequencing of the placenta (see Figure S4), serving as validation of the snRNAseq results. (**J**) Volcano plot illustrating DEGs identified in decidual cells.

### Single-nucleus RNA sequencing reveals cell type-specific vulnerabilities of the male placenta to MIA

While informative, bulk RNA-sequencing lacks the resolution to identify which components of the placenta are compromised following MIA. Thus, we performed single-nucleus RNA-sequencing (snRNAseq) to identify the placental cell types that may be particularly sensitive to poly(I:C). We mapped gene expression differences between placentas associated with poly(I:C)^mod^ versus male controls across 20 distinct cell types annotated based upon prior literature(*39, 42*). We grouped cell types into the following categories (genes enriched in each class are shown in brackets): (1) cells that make up the structural fabric of the placenta and support fetal health and growth (decidual cells [*Prl8a2*, *Cryab*], stromal cells [*Pbx1*, *Pgr*], two clusters of SpTs [*Taf7l*], glycogen cells [*Plac8*, *Igfbp7*], two clusters of epithelial cells [*Mecom*, *Cubn*], and parietal endoderm cells [*Lama1*, *Lamb1*]); (2) precursors that give rise to these cell types (SpT precursors [*Prl2c3*], syncytiotrophoblast precursors [*Epha4*,low *Ctsq*],and junctional zone precursors [*Prune2*]); (3) cells associated with the vasculature (sinusoidal trophoblast giant cells [*Ctsq*, *Lepr*, *Nos1ap*], syncytiotrophoblasts I [*Tfrc*], syncytiotrophoblasts II [*Slc13a4*, *Synb*], erythrocytes [*Hbb-y*], mesenchymal [*Col1a1*, *Col1a2*], endothelial cells [*Pecam1*], and maternally derived veinous endothelial cells [*Hdac9*]), and (4) immune cells (one cluster of maternally derived Natural Killer (NK) cells [*Ptprc*, *Gzmd*, *Fcrl6*] and one cluster of fetally derived macrophages [*Afp*, *Ptprc*, *Zeb2*]; Fig. 3B). The final dataset includes 197,237 nuclei passing a stringent QC pipeline in which unhealthy or dying cells, doublets, and ambient RNA were removed (Fig. S6).

To shed light on large-scale changes occurring within the placenta following MIA, we first asked whether the cellular composition of saline-exposed male and poly(I:C)^mod^ placentas differed by calculating the proportionality of cells across all types in both conditions. We find that several populations of fetally-derived cells that make up the middle layer of the placenta and share a developmental origin (SpT precursors, two populations of SpTs, glycogen cells [GCs], and junctional zone precursors) are significantly more abundant in poly(I:C)^mod^ placentas versus controls (Fig. 3C). To validate this upregulation, we quantified trophoblast giant cells (TGCs), identified based on their large size, within both conditions. TGCs are a terminally differentiated cell type derived from SpT progenitors that make up the junctional zone, the physical boundary between the maternal and fetal placental compartments(*43*). As suggested by snRNAseq data, we find that TGCs are significantly more abundant in poly(I:C)^mod^ placentas than in controls (Fig. 3D,E). In contrast, other cell types, including immune cells as well as cells involved in nutrient exchange between maternal and fetal blood vessels (stromal cells, sinusoidal trophoblast giant cells [S-TGCs, syncytiotrophoblasts [SynTI], fetally derived Hofbauer cells, endothelial cells, mesenchymal cells, and parietal endoderm cells) are less abundant in poly(I:C)^mod^ placentas (Fig. 3C). This suggests that the fetally-derived SpT lineage is expanded in the placentas of mice that exhibit developmental abnormalities in response to MIA. Conversely, cells that mediate interactions between maternal and fetal blood vessels as well as stromal cells are present at lower levels following MIA, suggesting compromised nutrient exchange.

Next, we applied differential gene expression analysis to uncover transcriptomic changes elicited by MIA. For every cell type, we observed robust transcriptomic shifts meeting a highly stringent threshold of 4.0 absolute fold change and FDR < 0.05 between poly(I:C)^mod^ and saline-treated male placentas, suggesting multi-cellular disruption arising within 24 hours of a single dose of poly(I:C) (Fig. 3F). Interestingly, the majority of differentially expressed genes (DEGs) for most cell types were downregulated rather than upregulated by MIA, potentially suggesting a generalized blunting of cellular function. These data suggest that transcriptomic changes occur in most cell types within the placentas of a vulnerable subpopulation of male embryos within 24 hours of exposure to MIA.

### Transcriptional pathways induced by MIA reflect inflammation and antigenicity

We first asked how the maternal structural and immune cell populations respond to poly(I:C) exposure, beginning with maternally derived NK cells (immune_1) and stromal cells. Genes upregulated by poly(I:C) in NK cells are highly enriched for mediators of programmed cell death. In particular, these cells upregulate several genes in the granzyme family of apoptosis-inducing secreted enzymes (e.g. *Gzmf*, *Gzmg*, *Gzmd*, *Gzme*, and *Gzmc*) which play an important role in the removal of damaged cells in response to inflammation, and have been previously associated with MIA(33). Concurrently, in response to poly(I:C), stromal cells dampen expression of genes associated with pregnancy, such as prolactin-like hormones (e.g. *Prl3b1*, *Prl7d1*, *Prl2c5*, *Prl7a1*, *Prl2c3*, and *Prl2c2*), endocrine signals that are necessary for promoting maternal immunotolerance of the fetus (Fig. 3G)(*44*). These stromal cells significantly blunt prolactin-like hormone signaling in response to MIA, while upregulating pro-inflammatory myeloid cell-related genes such as the complement component C3 and other aspects of type two immunity. Stromal cells also induce factors involved in antigen processing and presentation (e.g. H2-K1, H2-Q7), suggesting that structural tissues at the maternal-fetal interface undergo severe inflammatory stress. Thus, both maternally derived NK cells and local stromal cells exhibit significant activation in the placentas of poly(I:C)^mod^ embryos.

In contrast to maternally derived immune cells, MIA did not induce cell death pathways in fetally-derived macrophages (immune_2), but rather induced genes involved in cell recruitment and migration. For example, genes induced in these cells by MIA include the chemotactic cues *Cxcl2* (also known as MIP-2a) and *Cxcl10* (also known as IP-10) which play key roles in recruiting immune cells to sites of inflammation. Furthermore, members of the classical complement cascade (e.g. *C3)* are induced in fetally-derived immune cells by poly(I:C). Conversely, genes involved in lysosomal function, such as the Cathepsin family, are decreased in fetally-derived macrophages along with a blunting of prolactin-like hormone signaling as observed in maternal immune cells as well (Fig. 3H,I). Thus, MIA leads to the induction of pro-inflammatory pathways in both maternally and fetally derived immune cells, with maternally derived cells inducing cell death via granzyme production and fetally derived cells recruiting granulocytes and other immune populations to the fetal placenta.

Following poly(I:C) exposure, maternally-derived decidual cells also upregulate genes associated with a strong immune response to MIA, similarly to maternally derived immune cells. These cells induce mediators of antigen processing and presentation (e.g. *H2-K1*, *H2-D1*, *H2-Q7*) following MIA (Fig. 3J). These results are consistent with poly(I:C) directly activating decidual cells as previously reported(*45*). In addition to mounting a potentially antigenic response, decidual cells upregulate factors such as *Aqp1* and *Serpine1* which promote angiogenesis, and dampen the expression of genes associated with cellular motility and migration (e.g. *Amot* and *Kitl*) (Fig. 3J). These data suggest that a switch from a pregnancy-supporting, immunosuppressive microenvironment toward an inflammatory transcriptomic state characterizes the responses of both maternally-and fetally-derived placental cells to MIA.

### Genes encoding the extracellular matrix and immunosuppressive hormones are decreased in spongiotrophoblasts following MIA

Bulk RNA-sequencing suggests that the ECM is significantly compromised in the placentas of poly(I:C)^mod^ mice (Fig. S5), but the cell types contributing to this dysregulation are not clear. Our snRNAseq analysis addresses this question, revealing that SpTs are a primary cell type contributing to the loss of the ECM in the context of MIA. Specifically, SpTs decrease expression of several collagen (e.g. *Col1a1*, *Col1a2*, *Col12a1*) and laminin (e.g. *Lama2*, *Lama3*, and *Lama4*) family members following MIA, consistent with the results of our bulk RNA-seq analysis (Fig. 4A,B).

**Figure 4.**
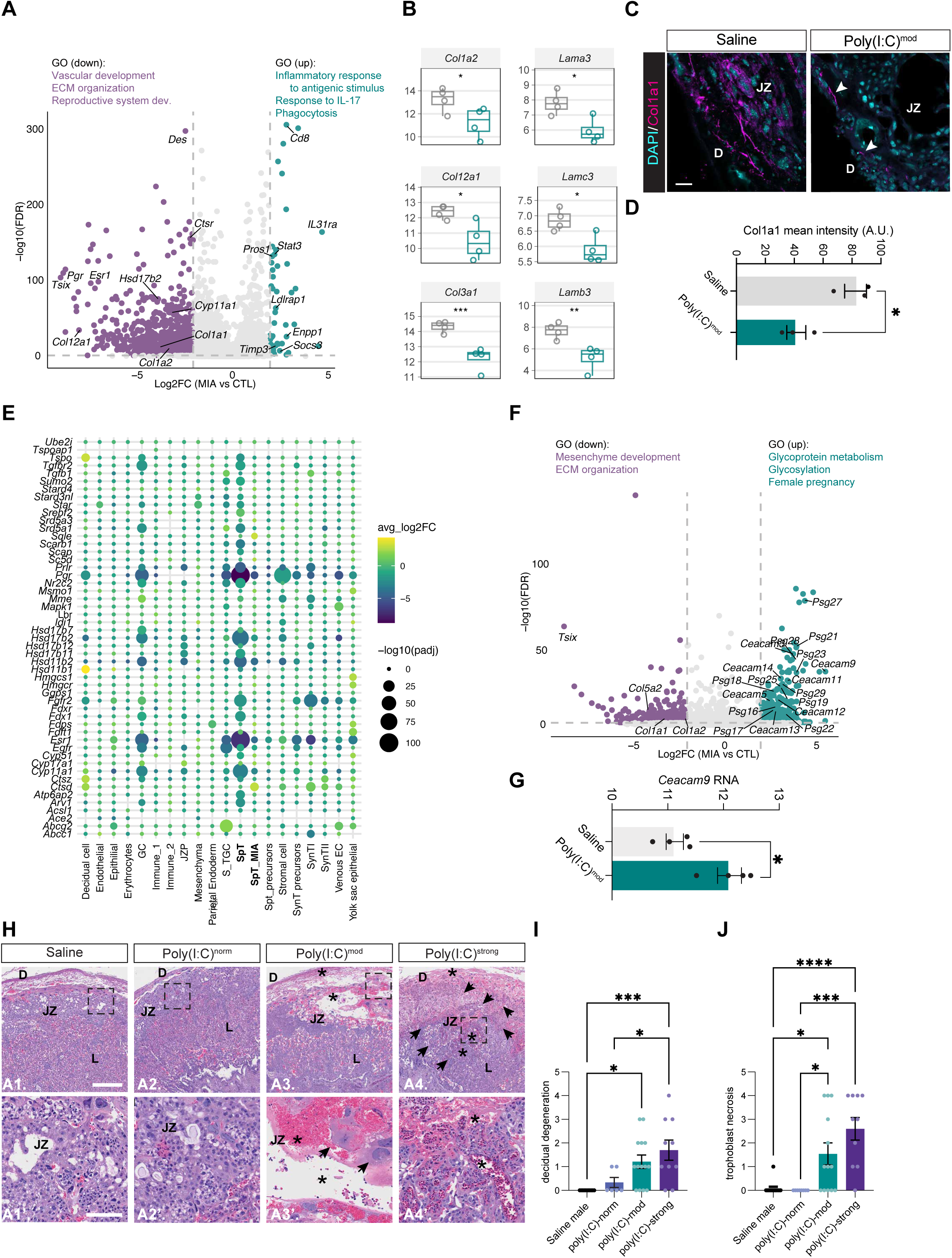
MIA reduces hormone and extracellular matrix transcripts in spongiotrophoblasts and induces a structural breakdown of the placental junctional zone. (**A**) Volcano plot illustrating DEGs identified in the largest cluster of spongiotrophoblasts (SpTs). (**B**) RNA levels of each given gene as measured by bulk RNA-sequencing of the placenta, serving as validation of the snRNAseq results. (**C**) Confocal images of placental sections from saline and poly(I:C)^mod^ male embryos immunostained for the collagen family member Col1a1 (magenta), DAPI shown in cyan. D, decidua; JZ, junctional zone; L, labyrinth. Arrows, examples of collagen deposition. Scale bar, 100 µm. (**D**) Quantification of the intensity of Col1a1 signal (A.U.) in placentas from saline and poly(I:C)^mod^ male embryos. Unpaired t test, *p < 0.05; n = 3 placentas per condition. (**E**) Dot plot illustrating changes in genes involved in hormone biosynthesis, signaling, and detection following MIA based upon snRNAseq. Fold change and statistical significance scales given on the right. (**F**) Volcano plot illustrating DEGs identified in SpT_MIA_ cells. (**G**) Levels of *Ceacam9* RNA in saline and poly(I:C)^mod^ placentas based upon bulk RNA-sequencing. Unpaired t test, *p < 0.05; n = 4 placentas per condition. (**H**) Microscopy images of H&E-stained placentas from male mice across conditions and phenotypes. D, decidua; JZ, junctional zone; L, labyrinth. Arrows, necrosis or degeneration. Asterisks, hemorrhagic congestion. Scale bar, 500 µm. Inset scale bar, 100 µm. Images in (H) also shown in Figure S8. (**I**),(**J**) Pathological scoring of degenerative signatures in male placentas based upon H&E staining. (**I**) Decidual degeneration. (**J**) trophoblast necrosis. (**I**),(**J**) One-way ANOVA with Tukey’s post test. *p < 0.05; ***p < 0.001; and ****p < 0.0001. For (**I**), n = 11 saline and 6 poly(I:C)^norm^, 14 poly(I:C)^mod^, and 10 poly(I:C)^strong^. For (**J**), n = 13 saline, 8 poly(I:C)^norm^, 13 poly(I:C)^mod^, and 10 poly(I:C)^strong^.

To validate this observation at the protein level, we performed immunofluorescence for Col1a1 in the placentas of males from control and poly(I:C)-treated dams representing all phenotypes: poly(I:C)^norm^ (no deficits), poly(I:C)^mod^ (moderate deficits), and poly(I:C)^strong^ (extreme deficits that can progress to reabsorption). We observe significant decreases in collagen deposition in male embryos from poly(I:C)^mod^ and poly(I:C)^strong^ groups, while collagen levels were intact in poly(I:C)^norm^ embryos which lacked developmental deficits (Fig. 4C,D). Thus, the placental ECM is dramatically compromised following MIA, but only in male embryos that exhibit developmental deficits, suggesting a direct link between the transcriptomic changes we observe in the placenta and the severity of the developmental abnormalities observed in the fetuses.

In addition to a decrease in ECM components, in response to MIA, SpTs joined several other cell types in exhibiting significantly decreased expression of transcripts involved in hormone biosynthesis and signaling (Fig. 4A,E). Whereas in immune cells most of the hormone-associated gene changes involve a dampening of prolactin-like hormones, SpTs strongly decrease their expression of *Hsd17b2* and *Cyp11a1*, both of which are involved in the biosynthesis of progesterone and testosterone, by over 8-fold. MIA also led to robust decreases in the expression of the progesterone receptor (*Pgr*) and estrogen receptor alpha (*Esr1*), which might be a compensatory response to the lower abundance of these hormones in general (Fig. 4A,E). Altogether, these data suggest that the dampening of ECM production and hormone signaling in fetally-derived SpT-lineage cells contributes to a loss of placental integrity at the junctional zone.

### A subpopulation of placental SpTs expressing high levels of glycoproteins is expanded in MIA

Consistent with SpTs being highly sensitive to poly(I:C) exposure, a subpopulation of SpTs, which we term SpT_MIA_ cells, is expanded in the placentas of poly(I:C)^mod^ embryos. These cells, which cluster distinctly from the largest SpT cluster described above, are twice as abundant in poly(I:C)^mod^ placentas as controls (Fig. 3B,C). Furthermore, differential gene expression analysis revealed that SpT_MIA_ cells induce remarkably high levels of two related classes of Ig-superfamily glycoproteins following MIA: pregnancy-specific glycoproteins (Psgs; *Psg27*,-*28*, - *21*,-*29*,-*19*,-*23*,-*25*,-*18*,-*17*,-*16*,-*22*, and-*26*) and Carcinoembryonic antigen-related cell adhesion molecules (Ceacams; *Ceacam3*,-*11*,-*5*,-*9*,-*14*,-*12*,-*13*, and-*1*)(Fig. 4F,G). While not widely studied in mice, work on the human placenta suggests that both classes of proteins can suppress maternal inflammation to protect the fetus during gestation(*46–49*). Interestingly, the two SpT populations in our dataset (SpT and SpT_MIA_) differ in their transcriptomic responses to MIA, as the largest cluster of SpTs decreases *Psg23* expression with only the smaller population of SpT_MIA_ cells increasing it. In line with this result, multiplexed fluorescence *in situ* hybridization for the SpT marker *Ascl2* and *Psg23* in placental sections revealed a lower percentage of *Psg23*-expressing SpTs in poly(I:C)^mod^ versus control males (Fig. S7A,B). These results implicate Psgs and Ceacams in the placental response to MIA, although whether these factors contribute to developmental deficits or, conversely, whether they represent an early protective response is not yet known.

### Human trophoblasts exposed to COVID-19 share antigenic responses and decreased collagen biosynthesis with MIA-exposed SpTs

To explore the potential translational relevance of our transcriptomic findings, we reanalyzed a single-cell RNA-sequencing dataset of placental tissue harvested at term from women with confirmed COVID-19 infections during pregnancy(*50*). That study reports inflammation within the placentas of COVID-19-infected women which was absent from healthy controls, and shows upregulation of granzymes *GZMA* and *GZMB*, consistent with our observation of the induction of granzymes in maternally derived immune cells following MIA (Fig. 3G). We next investigated the transcripts differentially expressed between COVID-19 and control conditions within villous cytotrophoblasts (VCTs) and extravillous trophoblasts (EVTs), two fetally-derived trophoblast populations involved in maternal-fetal interactions(*38*). We identified subsets of genes upregulated in both COVID-19 and poly(I:C)^mod^ (spongio)trophoblasts, indicating that the placental response to viral infection in humans and mice partially converges. As expected, gene ontology (GO) analysis revealed that transcripts upregulated in poly(I:C)^mod^ SpTs (our data), COVID-19-exposed VCTs, and COVID-19 exposed EVTs share functions in neural tube development, placental morphogenesis, and *in utero* embryonic development (Fig. S7C). Conversely, genes upregulated in both SpTs and VCTs (but not EVTs) are associated with myeloid cell activation and hemopoiesis, consistent with our observation that SpTs express inflammatory factors following MIA. SpTs and EVTs (but not VCTs) upregulate genes associated with T cell activation and proliferation and leukocyte cell:cell adhesion, in line with our observation that responses of SpTs to MIA reflect not only innate immune processes but also adaptive immunity. Concurrently, both SpTs exposed to MIA and EVTs exposed to COVID-19 downregulate genes associated with connective tissue development and collagen biosynthesis and metabolism (Fig. S7C). Together, these findings suggest that MIA in both humans and mice promotes inflammatory responses in placental trophoblasts concurrent with a reduction in components of the extracellular matrix.

### MIA disrupts the structural integrity of the decidua and junctional zone in poly(I:C)^mod^ and poly(I:C)^strong^ placentas

The dampening of ECM production and hormone signaling within placental cells suggests that MIA leads to a loss of structural integrity within the placenta. To test this hypothesis, we assessed placental architecture by scoring the severity of lesions revealed by H&E staining. We observed lesions that were restricted to the decidua and the junctional zone—characterized by classical pathological signs of hemorrhage, degeneration, congestion, and necrosis—in poly(I:C)^mod^ and poly(I:C)^strong^ placentas but not the placentas of male embryos that develop normally following MIA, or in the placentas of females from either treatment group (Fig. 4H-J, Fig. S8A-E, and Fig. S9). Given these striking sex-based differences, we also compared the architecture of untreated male and female placentas at E12.5, which indicated no lesions or other architectural differences between male and female placentas at baseline (Fig. S8F). Thus, a combination of bulk and single-cell transcriptomics, ECM immunostaining, and anatomical analyses reveals a male-specific breakdown in placental integrity downstream of MIA, particularly at the junctional zone, where maternal and fetal placental compartments meet. We further show that these disruptions only occur in the placentas of embryos exhibiting developmental abnormalities downstream of MIA.

### Innate and adaptive immune cells infiltrate the amniotic fluid following MIA

Each mouse embryo is contained in a unique amniotic sac, surrounded by amniotic fluid (AF), a liquid comprised of both fetally and maternally derived cells and molecules that undergo active exchange with the fetus. Given the structural deterioration of the junctional zone and the inflammatory responses of both maternally-and fetally-derived cells within poly(I:C)^mod^ placentas, we hypothesized that MIA alters the immune landscape of the AF, thereby contributing to developmental dysfunction in the embryo. However, while inflammatory signatures within the placenta have been observed in MIA models(*17, 25–27*), the impact of MIA on the composition of the AF has not been systematically investigated.

To measure the abundance and activation states of major immune cell classes within the AF, we subjected AF of male embryos across phenotypes and treatment groups to multispectral flow cytometry. This powerful approach allowed us to map the immune cell compartment of the AF across siblings within the same litters, obtaining >10,000 live cells per sample across the major immune lineages (defining markers given in parentheses): NK cells (NK1.1⁺/NKp46⁺), monocytes/macrophages (CD11b⁺), dendritic cells (CD11c⁺MHC-II⁺), T cells (CD3⁺), and B cells (CD19⁺; Fig. S10). In addition to assessing the abundance of immune cells, we also measured their activation states as detailed below.

Comparing AF composition between saline and poly(I:C)-treated male embryos across phenotypes revealed a significant expansion of several immune cell populations following MIA. For example, MIA induced a robust increase in the abundance of monocytes in the AF of poly(I:C)-exposed male embryos versus control males regardless of phenotype (Fig. 5A,B). Monocyte (including macrophage) activation as measured by CD45+,CD11b+, Ly6C+ co-expression trends toward higher levels in poly(I:C)^norm^ and poly(I:C)^mod^ embryos and is significantly higher in poly(I:C)^strong^ embryos versus saline, suggesting that the increased number of activated macrophages in the AF contributes to adverse phenotypes in the offspring. uNK cells are also significantly more abundant in the AF of poly(I:C)^mod^ embryos compared to saline, (Fig. 5A,C). These cells are derived from the maternal decidua, in line with our observation that uNK cell abundance within the placenta is decreased by MIA (Fig. 3B). Interestingly, dendritic cell numbers are decreased in the AF of poly(I:C)-exposed males compared to saline, again regardless of phenotypic severity (Fig. 5D). In terms of adaptive immunity, significant changes in the abundance of B cells were not identified, and numbers of B cells in the AF are low in general across conditions (Fig. 5A). However, somewhat unexpectedly, MIA significantly increases numbers of T cells within the AF of poly(I:C)^mod^ and poly(I:C)^strong^ placentas but not poly(I:C)^norm^ placentas, suggesting that, while changes in the abundance of innate immune cells within the AF extend across male embryos of all phenotypes, T cells are only present in the AF of embryos that exhibit deficits (Fig. 5A,E). Remarkably, the increased number of T cells occurs within poly(I:C)^mod^ and poly(I:C)^strong^ samples but not normally developing males even when all phenotypes were present in the same litter (Fig. 5F). Together, these findings provide evidence that MIA drives an influx of both innate and adaptive immune cells into the AF of male embryos within a 24-hour timeframe, poising maternally derived inflammatory signals to derail fetal development.

**Figure 5.**
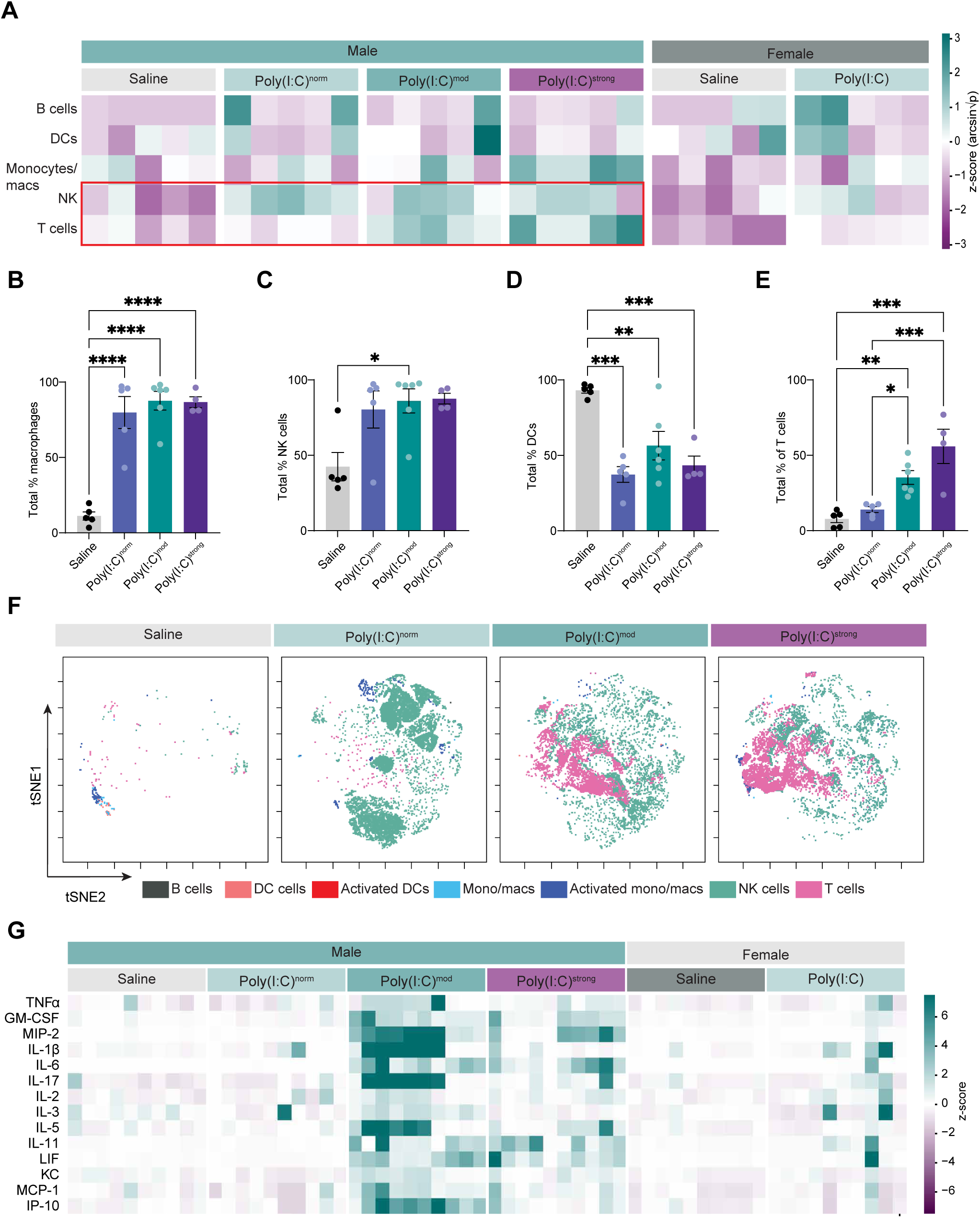
Innate and adaptive immune cells and pro-inflammatory cytokines infiltrate the amniotic fluid following MIA. (**A**) Heatmap demonstrating the abundance of distinct populations of immune cells in the amniotic fluid (AF) of individual embryos across sexes, conditions, and phenotypes. Rows, cell types given on left of heatmap. Columns, sex and condition. Scale on right. Red box emphasizes robust changes in NK and T cells in affected male placentas. (**B**)-(**E**) Quantification of the relative abundance (percentage of parental gate) of major immune cell populations in the AF: (**B**) monocytes/macrophages; (**C**) Natural killer (NK) cells; (**D**) Dendritic cells (DCs); and (**E**) T cells. For (**B**)-(**E**) One-way ANOVA with Tukey’s post test; *p < 0.05; **p < 0.01; ***p < 0.001; and ****p < 0.0001; n = 5 saline and poly(I:C)^norm^, n = 6 poly(I:C)^mod^, and n = 4 poly(I:C)^strong^ embryos collected from 5 litters per condition. (**F**) tSNE maps of immune cell abundance across saline and poly(I:C) conditions plotted by phenotype. Each map was derived from a single embryo and poly(I:C)-treated conditions were present in the same litter. (**G**) Heatmap demonstrating the induction of pro-inflammatory cytokines in the AF following MIA, scale shown on right.

### MIA induces a pro-inflammatory environment within the AF of poly(I:C)^mod^ and poly(I:C)^strong^ embryos

The remodeling of the immune cell repertoire in the AF of poly(I:C)-exposed male embryos provides evidence that inflammatory factors initiated in the dam lead to robust changes not just in the placenta but also in the direct environment of the fetus. To further delineate the immunological changes elicited by MIA in the AF and to relate these changes to developmental susceptibility, we performed multiplexed ELISAs to assess the expression of 37 cytokines and related soluble factors in females and males of all conditions and phenotypes. While most cytokines are unaffected by MIA regardless of phenotype, 14 cytokines exhibit an MIA-induced increase in expression within the AF of male fetuses exhibiting developmental abnormalities (i.e. poly(I:C)^strong^ and/or poly(I:C)^mod^ embryos) but not in normally developing males (poly(I:C)^norm^ mice; Fig. 5G). Strikingly, these 14 cytokines also fail to be significantly induced by poly(I:C) in the AF of female embryos (Table S1), suggesting a link to the developmental deficits observed in poly(I:C)^strong^ and/or poly(I:C)^mod^ males.

Cytokines induced by MIA in the AF of affected male embryos include the pro-inflammatory factors TNFα, Gm-CSF, MIP-2, IL-1β, IL-6, and IL-17. Both IL-6 and IL-17 have been previously associated with behavioral impairments in offspring of poly(I:C)-treated mice(*22, 24, 51–53*). In addition to this cohort of classical pro-inflammatory cytokines, several other Interleukin family members are induced by MIA in the AF of poly(I:C)^mod^ but not poly(I:C)^norm^ embryos: IL-2,-3,-5, and-11. As interleukins play a key role in mediating T and B cell activation, their selective presence within the AF of poly(I:C)^mod^ embryos further supports the idea that a transition from innate to adaptive immunity occurs in the direct environment of those fetuses (although at this early timepoint the innate immune response predominates), mirroring the responses observed in maternally-derived immune cells. Finally, LIF, KC, MCP-1, and IP-10, all of which are involved in immune cell chemotaxis and activation, also exhibit selective induction in poly(I:C)^mod^ versus poly(I:C)^norm^ embryos. Notably, increased levels of several of these cytokines (e.g. TNFα, IL-6, IL-17, IL-1β, and MCP-1) have been observed in the blood of mothers to children with ASD and the brains, cerebrospinal fluid, and blood of individuals with ASD, supporting the relevance of our findings to ASD in humans(*54–60*). Altogether, our analysis of the immune compartment of the AF revealed changes in cellular abundance and activation states that occur following MIA in all male embryos regardless of phenotype (with the exception of T cells which were absent from poly(I:C)^norm^ embryos), and changes in cytokine abundance that are selective to male embryos that are deleteriously affected by MIA. Similarly, female embryos do not exhibit an increase in cytokines within the AF following MIA (Table S1), suggesting that these cytokines contribute to the developmental deficits observed only in male embryos.

### IL-6 is required for MIA-driven deficits in male fetal development

While our data identify multiple MIA-induced cytokines within the AF, IL-6 is among the most highly induced in poly(I:C)^mod^ and poly(I:C)^strong^, but not poly(I:C)^norm^ or female, placentas (Fig. 6A,B). Moreover, IL-6 has been shown to promote ASD-like behavioral phenotypes of MIA_poly(I:C)_ offspring(*24, 52*). Thus, we hypothesized that IL-6 links the early developmental deficits that we describe here to the ASD-like behavioral impairments previously characterized in adult offspring of MIA_poly(I:C)_ mice.

**Figure 6.**
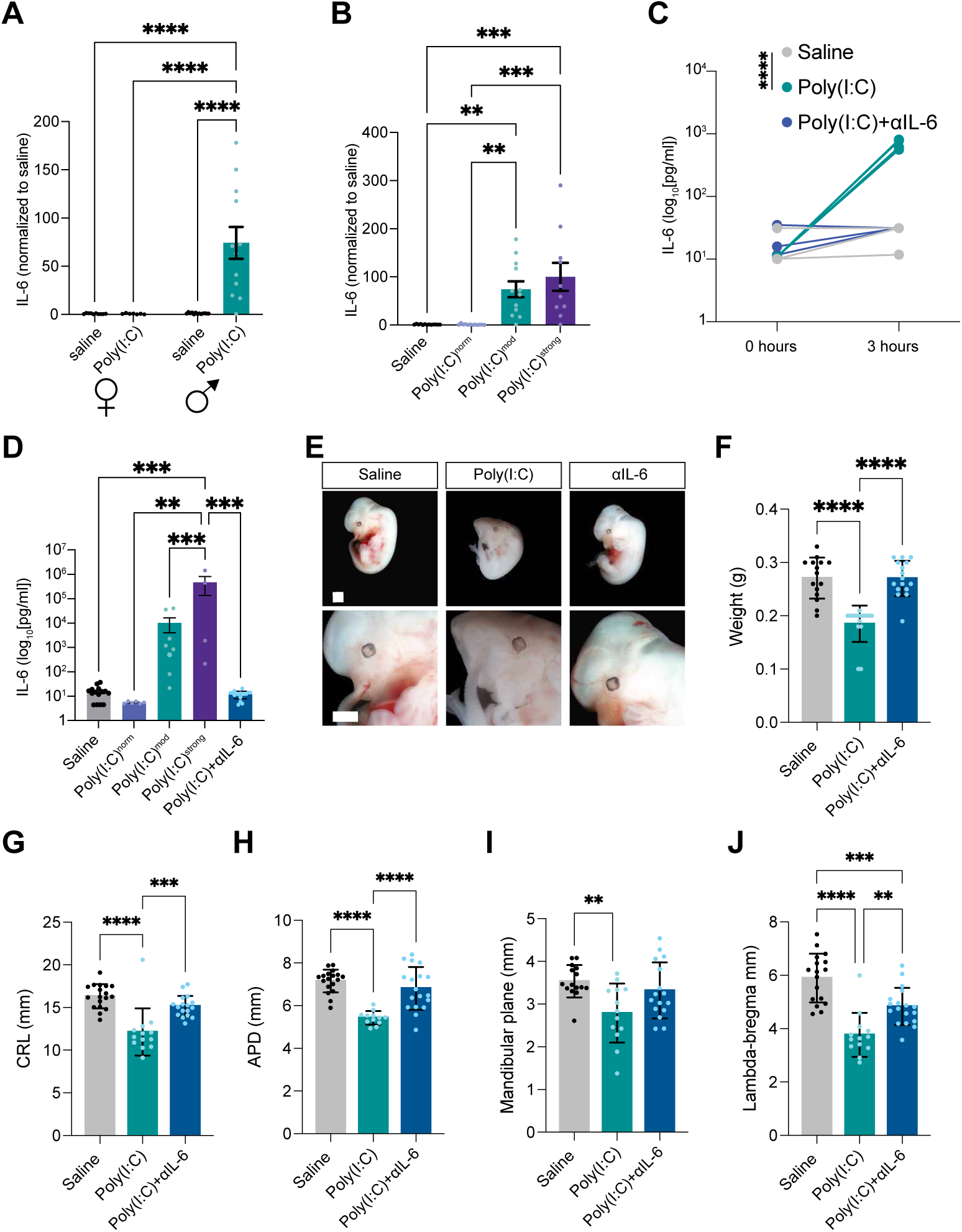
IL-6 is required for developmental abnormalities in male embryos. (**A**) IL-6 protein levels in the AF of male and female embryos from saline and poly(I:C) treatment groups. Two-Way ANOVA with Tukey’s post test, ****p < 0.0001; n = 10 saline female, 13 saline male, 6 poly(I:C) female, and 13 poly(I:C) male embryos. (**B**) IL-6 levels measured across male embryos of each phenotype. One-Way ANOVA with Tukey’s post test; **p < 0.01; ***p < 0.001; n = 13 saline, 12 poly(I:C)^norm^, 13 poly(I:C)^mod^, and 10 poly(I:C)^strong^ embryos. (**C**) Levels of IL-6 in the serum of dams exposed to poly(I:C) (teal), poly(I:C) and the anti-IL-6 neutralizing antibody MP5-20F3 (αIL-6, indigo), or saline (gray) measured by ELISA. Unpaired t test, ****p < 0.0001; n = 4 litters/condition. (**D**) Analysis of IL-6 in the AF, demonstrating depletion of IL-6 in the AF of αIL-6-exposed embryos. One-way ANOVA with Tukey’s post test; **p < 0.01; ***p < 0.001; n = 15 saline, 6 poly(I:C)^norm^, 8 poly[I:C]^mod^, 4 poly[I:C]^strong^, and 18 (αIL-6 + poly(I:C). (**E**) Example images of fetuses from each treatment group at E13.5. Scale bars, 1 mm. (**F**)-(**J**) Measurement of fetal morphology across conditions. Parameters measured: (**F**) fetal weight (g); (**G**) Crown-rump length (mm); (**H**) Anterior-posterior diameter (mm); (**I**) Mandibular plane length (mm); and (**J**) lambda-bregma distance (mm). One-way ANOVAs with Tukey’s post test, **p < 0.01; ***p < 0.001; ****p < 0.0001. For (**F**), n = 15 saline, 16 poly(I:C), and 16 (αIL-6 + poly(I:C) embryos. For (**G**), n = 18 saline, 13 poly(I:C), and 18 (αIL-6+ poly(I:C) embryos. For (**H**), n = 18 saline, 12 poly(I:C), and 18 (αIL-6+ polyI:C) embryos. For (**I**), n = 16 saline, 13 poly(I:C), and 16 (αIL-6+ poly(I:C)). For (**J**), n = 17 saline, 13 poly(I:C), and 18 (αIL-6+ poly(I:C)) embryos.

To test this hypothesis, we used the MIA_poly(I:C)_ paradigm but included neutralization of maternal IL-6 via intraperitoneal injection of the IL-6 neutralizing antibody MP5-20F3 one hour prior to exposure to poly(I:C). We validated depletion of IL-6 by ELISA of maternal blood as well as the AF (Fig. 6C,D). We find that, while poly(I:C) still elicits developmental abnormalities and structural defects in about 30% of embryos and their corresponding placentas, respectively, at E13.5, it does not affect fetal or placental integrity when IL-6 is neutralized (Fig. 6E-J and Fig. S11). Thus, maternal IL-6 is not only necessary for the appearance of ASD-like behaviors in adult offspring following MIA, it is also necessary for the abnormalities in fetal development that emerge within 24 hours of poly(I:C) exposure. This finding suggests that IL-6 links inflammation at the maternal-fetal interface to the behavioral deficits that emerge in male fetuses following exposure to MIA.

## DISCUSSION

While the vast majority of mechanistic studies on the biological drivers of ASD focus on the brain, the strong association between MIA and ASD risk indicates that prenatal immunological dysfunction is a critical, but understudied, contributor to disease etiology. By characterizing changes in fetal health that occur within 24 hours of poly(I:C) exposure, we identify male-specific developmental abnormalities that emerge following MIA. This result is consistent with published evidence that male offspring of MIA_poly(I:C)_ mice exhibit more extreme behavioral and cognitive symptoms than females(*61, 62*), as well as emerging evidence that males exhibit heterogeneity in their behavioral responses to MIA(*9, 12, 36, 37, 63*). Our data suggest that, in cases where inflammation is involved, male bias in ASD arises in gestation rather than or in addition to accumulating over postnatal development.

At the mechanistic level, our analyses identify the placenta—particularly fetally derived SpTs at the junctional zone—as a primary target of MIA. snRNAseq and histological analyses reveal three convergent effects of maternal inflammation: (1) induction of immune and antigen-response programs across maternal NK cells, fetal macrophages, and stromal cells; (2) suppression of extracellular matrix and endocrine pathways, which are essential for maintaining an immunosuppressive placental microenvironment, in SpTs; and (3) structural degradation of the junctional zone, coinciding with immune cell and cytokine infiltration into the AF. Finally, we show that IL-6 infiltrates the AF and is required for the developmental deficits observed in male embryos exposed to MIA, consistent with IL-6 being necessary for the adult offspring of MIA mice to exhibit cognitive and behavioral deficits in the MIA_poly(I:C)_ model(*52*). Thus, our work demonstrates that MIA induces a switch from an immunosuppressive placental microenvironment toward an inflammatory state characterized by the expression of factors that regulate the host immune response to non-self entities (Fig. S12).

Based on these results, we hypothesize that two factors are necessary to give rise to developmental abnormalities following MIA: (1) an abundance of non-self proteins, i.e. antigens, derived from the fetus, and (2) an inflammatory placental microenvironment. In a healthy pregnancy, although male fetuses express high levels of paternally derived proteins, the placental microenvironment suppresses inflammation to protect and support the embryo. However, in the context of MIA, the placental microenvironment loses its immunosuppressive properties, allowing these fetally-derived antigens to activate maternal immune cells which ultimately injures the placenta and compromises fetal health. Although the placentas of female embryos exposed to poly(I:C) are also likely to lose some immunosuppressive properties as they are exposed to the same maternal factors, the lower burden of paternally derived antigens allows the placenta to maintain healthy development of the female fetus.

Although our study is among the first to provide an unbiased analysis of changes at the maternal-fetal interface that are elicited within 24 hours of MIA, our results are largely consistent with prior studies interrogating cellular and molecular changes resulting from midgestational exposure to poly(I:C). In particular, placental levels of IL-6 and TNFα, inflammatory cytokines that we find to selectively accumulate within the AF of poly(I:C)^mod^ embryos, were significantly upregulated within the first 24 hours of poly(I:C) exposure(*24, 26, 64*). Concurrently, this upregulated IL-6 was shown to elicit JAK/STAT3 signaling within the SpT layer of the placenta, consistent with our result that SpTs undergo transcriptomic changes in response to MIA that contribute to a loss of structural integrity at the junctional zone. Studies interrogating MIA-driven changes within the placenta at later timepoints following MIA (i.e. at E15.5, E17, and E18.5) demonstrate that at least some of the inflammatory signatures we identified at E13.5 persist across gestation. Specifically, at E15.5 (72 hrs after poly(I:C) exposure), Th17-related inflammatory signaling was strongly upregulated within the placenta, in line with our observation that IL-17-related gene expression is upregulated in placental immune cells by E13.5(*65*). When placentas were analyzed closer to gestation, increased levels of cell death were found within the junctional zone, consistent with our placental pathology data and supporting the possibility that we may be able to identify some of these affected male mice at a later stage of development(*66*). Thus, although Monteiro et al (66) did not characterize the structural breakdown of the placenta with the same granularity as applied in our study (and, given that their analysis was performed several days later than ours, the difference could be based upon timepoint analyzed), evidence of cell death at the junctional zone is in line with our observations. Other groups have shown inflammation to persist at E18.5, where numbers of placental Tregs are decreased (consistent with our single-cell sequencing result that some immune cells are less abundant within the placenta following MIA)(*25*). Macrophages were also shown to be important for MIA-driven behavioral phenotypes as these phenotypes were abolished by macrophage depletion, in line with our finding that macrophages accumulate within the AF of poly(I:C)^mod^ embryos(*25*). Finally, it was shown that Gm-CSF, IL-6, and TNFα, three factors that also accumulate within the amniotic fluid of poly(I:C)^mod^ embryos, were upregulated only in male placentas at E17.5, consistent with the male-specific effects observed in our study(*17*). Thus, our work both confirms prior results and significantly expands our understanding of how inflammation at the maternal-fetal interface contributes to embryonic developmental dysfunction in the context of MIA.

Our results are consistent with an immunological basis of some cases of ASD. In line with this interpretation, individuals with autoimmune disorders are more likely to have offspring diagnosed with ASD (*67–70*). Likewise, the increased abundance of pro-inflammatory cytokines within the blood of mothers to children with ASD and the brains, cerebrospinal fluid, and blood of individuals with ASD point toward a chronic inflammatory component to the pathophysiology of the disorder in a subset of patients, particularly in males. Notably, these cytokines include TNFα, IL-6, IL-17, and IL-1 (*55–60, 71, 72*) which were also identified in our data (although it should be noted that a recent study failed to observe the upregulation of some of these cytokines in the fetal brain following MIA(*73*)). IL-6 may be of particular relevance therapeutically as our data indicate that IL-6 is required for the emergence of MIA-driven developmental deficits, while work from other labs suggests that IL-6 is also necessary (and sufficient) to promote ASD-like behaviors in adult offspring exposed to MIA(*51, 52*).

Other evidence in humans points toward a potential role for maternal-fetal antigenicity in MIA-driven neurodevelopmental dysfunction as well. For example, maternal antibodies against fetal brain proteins were observed in about 12% of parents of children with ASD, and higher levels of these antibodies were positively correlated with the severity of neurodevelopmental impairments in autistic offspring(*74–78*). Moreover, ASD is more common in first-born children, potentially because this pregnancy is the first introduction of the maternal immune system to the fetus’s antigens(*79*). ASD is also more common in children born to an older father(*79, 80*). This is relevant because advanced paternal age is associated with increased mutations in the sperm(*81, 82*), which could create new fetal antigens against which the maternal immune system may mount a response. Finally, although the question of whether maternal-fetal antigenicity contributes to male bias in ASD by rendering males more susceptible to maternal-fetal immune conflict remains unresolved, this immune incompatibility is already thought to contribute to other embryonic problems such as pre-eclampsia and intrauterine demise(*83*). Of note, our poly(I:C)^strong^ cohort appears to be in the process of intrauterine demise within 24 hours of MIA, consistent with this association. These observations suggest that some cases of ASD may share pathophysiological mechanisms, but have less extreme outcomes, with conditions like pre-eclampsia.

### Limitations of the study

This study has several limitations. The use of a mouse model limits direct extrapolation to human pregnancy, and aspects of placental architecture and immune regulation differ between the species. Experiments expanded to crosses between unrelated (i.e. not inbred) mouse strains could be an informative future direction, as these would be more likely to unveil antigenicity between mother and fetus given their distinctive genetic backgrounds. Such crosses may reveal not only increased susceptibility of males to MIA, but could shed light on factors influencing female susceptibility as well. Another caveat that is important to consider is the high degree of variability that hampers MIA models, wherein factors as basic as the commercial source of the poly(I:C) can dramatically change the impact of MIA on fetal neurodevelopment(*32–34*). Thus, it is possible that some of the effects of poly(I:C) exposure identified here may not occur, or may be more subtle, in dams exposed to alternative sources of poly(I:C). In fact, this could be one reason that robust changes in placental integrity downstream of poly(I:C) have not been previously reported. Moreover, our finding that close proximity to other male fetuses in the womb increases risk of developmental abnormalities is interesting and may reflect, for example, an accumulative inflammatory effect that can spread from one male embryo, for example an embryo undergoing demise, to the other. However, whether this finding extends to the behavioral manifestation of ASD in adult offspring is not known and requires follow-up as a prior study showed that embryos neighbored by females were more likely to exhibit behavioral deficits following MIA(*37*). Finally, our focus on early embryonic outcomes precluded direct assessment of how cytokines in the AF influence fetal brain development. Despite these caveats, our findings define early determinants of susceptibility versus resilience to prenatal inflammation and highlight the maternal–fetal interface as a critical site of ASD risk.

## MATERIALS AND METHODS

### Animal models

All experiments were performed in compliance with protocols approved by the Institutional Animal Care and Use Committee (IACUC) at Cold Spring Harbor Laboratory. The following mouse lines were used in the study: C57Bl/6J (The Jackson Laboratory, JAX:000664) and BALB/cJ (JAX:000651). Mice were maintained in standard housing at 20–22°C with 40– 55% humidity on a 12-hour light/dark cycle, with food and water provided ad libitum.

### Maternal Immune Activation (MIA)

C57BL/6J or BALB/c mice maintained in-house for ≥3 generations were used for breeding unless otherwise indicated. Dams were bred between ages P60-P150 and male breeders were aged between P80-P200. Estrous cycle stage was determined via daily vaginal lavage, and females in proestrus were housed overnight with proven males. The visible presence of sperm or a vaginal plug the following morning marked embryonic day E0.5, and the male was removed at this time. MIA was induced at E12.5 (with smaller cohorts being induced at E9.5 or E14.5) by intraperitoneal injection of polyinosinic–polycytidylic acid [poly(I:C), 20 mg/kg; Sigma-Aldrich P9582, lots 2137379, 329448, and 373973)] or sterile saline as a control. While this source of poly(I:C) has been shown to exhibit batch-to-batch variability(*33, 34*), we observed similarly high rates of success in eliciting an IL-6 increase in maternal serum within three hours of MIA, and the effects of poly(I:C) on fetal development and the maternal-fetal interface were consistent across lots. Dams were euthanized 24 hours post-treatment (E13.5 for most experiments) and embryos, placentas, and amniotic fluid were collected.

### Validation of MIA

Maternal blood was collected immediately before poly(I:C) injection at 0 hours and 3 hours post-injection via facial or submandibular vein puncture using a sterile 5 mm lancet (Goldenrod Animal Lancet). Approximately 200 µL of whole blood was collected per time point, centrifuged at 1,400 × g for 10 minutes at 4°C, and ≥25 µL of serum was transferred to Protein LoBind microtubes and stored at 4°C for up to 24 hours. Maternal IL-6 concentrations were quantified using a commercial ELISA (R&D Systems DY406-05). Absorbance at 450 nm was measured, and concentrations were calculated by four-parameter logistic fitting. Dams that failed to mount a sufficient IL-6 response or exhibited severe illness/spontaneous abortion were excluded from further analyses.

### Quantification of developmental abnormalities in embryos

Embryos were imaged under a stereomicroscope Nikon SMZ1500, and morphological measurements were obtained using ImageJ. In addition to fetal weight, parameters measured include crown–rump length (CRL), anterior–posterior diameter (APD), hind limb length, lambda– bregma distance, mandibular plane length, and muzzle–auricular angle. In addition, the percentage of the surface of each embryo displaying vascular hemorrhaging was quantified. Additional categorical scoring captured abnormalities in the ears, whisker pad, tail, and spine. Measurements were summarized at the litter level to assess within-litter penetrance.

### MCA Analysis

Morphological traits were binarized and analyzed by Multiple Correspondence Analysis (MCA) implemented in Python using prince, pandas, and matplotlib packages. Embryos were assigned to poly(I:C)^norm^, poly(I:C)^mod^, or poly(I:C)^strong^ categories based on predefined morphological criteria. Group ellipses were derived from covariance matrices and scaled to encompass approximately 95% of observations, representing dispersion rather than confidence intervals. Ellipses were centered at group centroids and aligned to the principal direction of variance. All visualizations were exported as vector-based PDF files for downstream editing.

### Sex determination

Embryonic sex was determined by PCR genotyping of tail samples using primers for *Rbm31* which takes different forms when expressed on the X versus the Y chromosome. When the Y chromosome is present, a larger fragment is amplified, allowing us to confidently annotate embryos as male or female depending upon whether this band was present (males) or not (females). We further validated these sex assignments through PCR for the Y-chromosome-specific gene *SRY*. Primer sequences used were: *Rbm31* (forward), 5’-CACCTTAAGAACAAGCCAATACA-3’; *Rbm31* (reverse), 5’-GGCTTGTCCTGAAAACATTTGG-3’; *SRY* (forward), 5’-GGAGTGGCATTTTACAGCCTGC-3’; *SRY* (reverse), 5’-TGGAGTACAGGTGTGCAGCTCT-3’. PCR products were resolved by agarose gel electrophoresis.

### Bulk RNA sequencing

Placentas were collected from saline-treated control males or poly(I:C)^mod^ males under a Nikon SMZ1500 stereomicroscope and weighed. About 20 mg of placental tissue was then subjected to RNA extraction. Tissue was transferred to QIAGEN Buffer RLT supplemented with β-mercaptoethanol used to enhance RNase inactivation and homogenized with ceramic beads using a BeadBug™ 6 microtube homogenizer (Benchmark Scientific) until no visible fragments remained. Total RNA was purified using the RNeasy Mini Kit (QIAGEN, cat. no. 74106) according to the manufacturer’s instructions, including on-column DNase digestion using the RNase-Free DNase Set (QIAGEN; Buffer RDD). Purified RNA was eluted in RNase-free water and stored at −80°C until library preparation and sequencing. Libraries were sequenced on an Illumina NextSeq 2000 using paired-end 150-bp reads (300-cycle configuration), yielding ∼400 million reads in total across all samples.

Paired-end FASTQ files were initially assessed for sequencing quality using FastQC (Babraham Bioinformatics). Adapter trimming and quality filtering were performed using fastp in paired-end mode with a Q20 quality threshold and overrepresentation analysis enabled, followed by post-trimming quality assessment using FastQC. High-quality reads were aligned to the mouse reference genome (mm10) using the splice-aware aligner STAR. The STAR genome index was generated from the mm10 genome FASTA and corresponding gene annotation GTF obtained from a publicly available reference package (refdata-gex-mm10-2020-A). Alignments were performed with multithreading and on-the-fly decompression of gzipped FASTQ files, and outputs were generated as coordinate-sorted BAM files. Gene-level read counts were quantified from sorted BAM files using HTSeq-count, counting reads overlapping annotated exons and summarizing counts at the Ensembl gene_id level in unstranded mode.

Differential gene expression analysis was conducted in R (v4.4.3) using DESeq2. Raw gene count matrices were combined across samples and aligned with sample metadata, and differential expression was modeled using negative binomial generalized linear models with experimental condition specified as the primary explanatory variable. Resulting *P* values were adjusted for multiple hypothesis testing using the Benjamini–Hochberg false discovery rate correction. Gene identifiers were annotated by mapping Ensembl gene IDs to official gene symbols using biomaRt with the Ensembl *Mus musculus* database. For visualization, size-factor– normalized counts were log₂-transformed after adding a pseudocount of 1, and expression levels of selected genes were visualized using boxplots overlaid with individual sample points; statistical significance was annotated directly from adjusted *P* values (without re-testing). Data manipulation and visualization were performed using dplyr, tidyr, and ggplot2. Functional enrichment analyses were carried out using clusterProfiler, with curated gene sets obtained from msigdbr and enrichment testing performed using fgsea, as indicated. Heatmaps were generated using ComplexHeatmap or pheatmap, as specified.

### Single-nucleus RNA sequencing (snRNAseq)

#### Tissue preparation

For snRNAseq, placentas from saline-exposed and poly(I:C)^mod^ male embryos were flash-frozen in liquid nitrogen. Frozen tissues were thawed in 1 ml of high sucrose buffer containing 0.25 M sucrose, 25 mM KCl, 5 mM MgCl₂, 20 mM Tricine-KOH, pH 7.8, 0.15 mM spermine tetrahydrochloride, 0.5 mM spermidine trihydrochloride, 1 mM DTT in the presence of RNAse inhibitors. The samples were mechanically dissociated using a Dounce homogenizer containing IGEPAL CA-630 (Sigma-Aldrich). Nuclei were collected from the interface of 30% and 40% of iodixanol after 18 minutes of density gradient ultracentrifugation at 10,000 x g. Purified nuclei were counted using a hemocytometer and processed as per the 10x Chromium Next GEM Single Cell 3’ v3.1 kit to recover approximately 10,000 nuclei/ml. Libraries were sequenced at The Center for Applied Genomics (TCAG), The Hospital for Sick Children, Toronto, ON, Canada.

#### Bioinformatics workflow

snRNAseq data processing and downstream analyses were performed using the Cell Ranger and Seurat pipelines. Raw sequencing reads were demultiplexed and aligned to the mouse reference genome (mm10) using Cell Ranger (10x Genomics, v7.1.0), generating gene-by-nucleus count matrices for each library. Ambient RNA contamination, a common artifact of droplet-based sequencing platforms, was corrected using SoupX (v1.6.2), which automatically estimates and removes background RNA signatures originating from empty droplets. This correction improved cluster separation and biological signal clarity in subsequent analyses.

Following data import, nuclei with fewer than 200 or more than 2,500 detected genes, as well as those with greater than 5% mitochondrial gene content, were excluded to remove low-quality nuclei and potential doublets. Data normalization and variance stabilization were performed using Seurat’s SCTransform function to enable accurate scaling and gene expression comparisons across samples. For integrated analyses across biological replicates and experimental conditions, datasets were merged using the Seurat integration pipeline (v5.0.1) in conjunction with Harmony batch correction (v0.1.1), which mitigates technical variability while preserving biologically meaningful differences.

Dimensionality reduction was performed using principal component analysis (PCA), with the top 30 principal components retained for downstream analyses. A shared nearest-neighbor (SNN) graph was constructed based on these components to capture transcriptomic relationships among nuclei. Clustering was performed using the Louvain algorithm (resolution = 0.8), and low-dimensional visualization was generated using Uniform Manifold Approximation and Projection (UMAP) with 30 dimensions. Cluster identities were assigned based on the expression of canonical lineage-specific marker genes.

#### Differential gene expression (DGE) analysis

Differential gene expression analyses were performed for each identified cell type using the Model-based Analysis of Single-cell Transcriptomics (MAST) framework, implemented through Seurat’s FindMarkers function. Biological replicates were included as a latent variable to account for inter-sample variability within each experimental group, ensuring robust identification of consistent transcriptional changes. To restrict analyses to reliably expressed genes, only genes detected in at least 5% of nuclei in either condition were tested. Differentially expressed genes (DEGs) were defined by an FDR-adjusted p-value < 0.05 and an absolute log₂ fold change ≥ 4.

#### Gene Ontology (GO) Enrichment Analysis

Functional enrichment analysis was performed on significantly differentially expressed genes using the clusterProfiler package (v4.0). Gene sets included DEGs with FDR-adjusted p-values < 0.05 and log₂ fold changes ≥ 4 (upregulated) or ≤ −4 (downregulated). Gene Ontology (GO) terms containing fewer than 10 or more than 500 genes were excluded to avoid overly narrow or broadly defined categories. Multiple hypothesis testing was corrected using the Benjamini–Hochberg method, and GO terms were considered significantly enriched if they met both an FDR < 0.05 and a gene ratio > 0.1. Upregulated and downregulated gene sets were analyzed independently to identify condition-specific biological pathways.

### Single-molecule fluorescence in situ hybridization (smFISH)

Coronal sections (12 μm) of placenta were generated using a Leica CM3050S cryostat, collected on Superfrost Plus slides, and stored at −80 °C until use. Multiplexed smFISH was performed using the RNAscope platform (Advanced Cell Diagnostics [ACD], Biotechne) following the manufacturer’s instructions for fresh-frozen samples (kit v2). Probes targeting *Ascl2 (Mash2)* and *Psg23* transcripts were used. For imaging, 40X confocal images were acquired as 5×5 tile scans using an LSM 780 Zeiss microscope. Three mice per condition were analyzed, with two images collected per mouse.

To quantify the proportion of cells positive for *Ascl2* and *Psg23*, images were processed as follows. First, the DAPI channel was thresholded and binarized, then expanded using the dilate function. The expanded DAPI mask was subjected to a watershed filter to ensure separation of adjacent nuclei. This mask was used to generate cell-specific regions of interest (ROIs), each corresponding to a single cell. For each ROI, the mean gray value within the *Ascl2* and *Psg23* channels was measured. The resulting values were exported into Excel for intensity-based classification. Background intensity for each marker was manually defined, and ROIs with mean intensities above background were assigned a value of 1 (positive), whereas ROIs below background were assigned a value of 0 (negative). These binarized datasets were used to calculate the percentage of cells positive for each transcript and the proportion of cells co-expressing *Ascl2* and *Psg23*.

### Human scRNA-seq Re-analysis

The GSE171381 dataset represents single-cell RNA sequencing of placental tissue from mothers infected with COVID-19 during pregnancy. Raw count matrices and metadata from GSE171381 were imported into Seurat(v5). Ambient RNA contamination was removed using decontX (celda). Decontaminated counts were normalized and processed using standard Seurat workflows. Single-cell clustering was performed according to the original published cell-type annotations (“annotation_merged”). Differentially expressed genes between COVID-19 and control samples were identified within each cell type using Seurat’s Wilcoxon test and MAST. Only genes exceeding a log₂ fold change ≥ 2 with a p-value ≤ 0.05 were considered. Significant DEGs from human villous cytotrophoblasts (VCT), extravillous trophoblasts (EVT), and mouse spongiotrophoblasts (Spt) were compiled. Gene symbol formatting was standardized, and overlapping sets were quantified in Excel. GO enrichment analysis was performed on overlapping DEGs using clusterProfiler. Analyses included biological process (BP), cellular component (CC), and molecular function (MF) ontologies, and the top-enriched terms were visualized using standard plotting functions within the package.

### Placental pathology

Mouse placentas were harvested and immediately immersion-fixed in ice-cold 4% paraformaldehyde (PFA) in PBS overnight at 4°C. After fixation, tissues were transferred to 70% ethanol and processed for paraffin embedding. Paraffin sections were stained with hematoxylin and eosin (H&E) and cut parallel to the chorionic plate and evaluated by a placental pathologist blinded to condition. Degeneration, vascular congestion, trophoblast degeneration, and necrosis were scored on a 0–5 scale. Representative images were acquired under identical imaging settings. Scores were compared across groups by one-way ANOVA followed by Tukey’s post hoc correction, or two-way ANOVA where appropriate.

### Immunofluorescence

Placentas were harvested and immediately immersion-fixed in ice-cold 4% paraformaldehyde (PFA) in PBS overnight at 4°C. Following fixation, tissues were cryoprotected by incubation in 15% sucrose and then 30% sucrose (in PBS) until fully infiltrated, embedded in OCT (stored at −80°C), and sectioned into transverse sections perpendicular to the chorionic plate at 12 μm onto Superfrost Plus slides using a Leica CM3050 S cryostat. Sections were washed in PBS and blocked in blocking solution (PBS containing 5% normal goat serum [NGS] and 0.3% Triton X-100 [TX-100]) for 1 hour at room temperature, then incubated overnight at 4°C in primary antibody (Rabbit anti-Col1a1, Cell Signaling Technologies 72026S, [1:300], RRID AB_2904565) diluted in blocking solution. The next day, sections were washed 3 times for 10 minutes per wash in PBS before incubation in secondary antibody Alexafluor 555 anti-rabbit (Thermofisher A21428; [1:1000], RRID AB_2535849) diluted in probing solution (PBS with 5% normal goat serum and 0.3% Triton X-100) for 2 hours at room temperature. Sections were then washed in PBS, stained for 10 minutes with NucBlue^TM^ (Invitrogen,R37605) mounted in fluoromount-G (SouthernBiotech), and cover-slipped.

Col1a1 fluorescence was imaged on a Zeiss LSM 710 confocal microscope as Z-stacks using identical acquisition settings across groups and converted to maximum intensity projections in Fiji/ImageJ. Images were processed identically across samples, including rolling-ball background subtraction applied uniformly. Anatomically matched ROIs were defined with comparable quantified tissue area across samples, and Col1a1 signal was quantified as mean fluorescence intensity (mean gray value) within each ROI, excluding saturated pixels and tissue artifacts. Multiple ROIs/sections were averaged per biological replicate and exported for downstream statistical analysis.

### Flow cytometric analysis of amniotic fluid

Amniotic fluid was collected from individual embryos into pre-chilled low-bind microtubes and kept on ice. Samples were diluted in cold FACS buffer (PBS + 2% FBS) and passed through a 40–70 µm cell strainer to remove debris and aggregates. Cells were pelleted by centrifugation at 500 × g for 7 min at 4°C. Supernatants were removed and retained for cytokine measurements when indicated, and cell pellets were gently resuspended in FACS buffer. Cells were incubated with anti-mouse CD16/32 (Fc receptor-blocking antibody; BioLegend, clone 2.4G2) to prevent non-specific antibody binding for 10 min at 4°C, followed by staining with a fixable viability dye and surface antibody cocktails (panel in supplementary material S10A) for 25–30 min at 4°C in the dark. Samples were washed twice with FACS buffer and resuspended in FACS buffer for acquisition.

Single color reference controls were prepared using both cells and antibody capture beads, together with an unstained control, and acquired in parallel. Data were acquired on a Sony ID7000 spectral flow cytometer, and spectral unmixing, including autofluorescence extraction when applicable, was performed using the reference controls. Gating strategies were standardized across samples to define major immune lineages: monocytes/macrophages, NK cells, dendritic cells, and T cells. Dimensionality reduction was performed using viSNE on the Cytobank platform. Antibodies used in flow cytometry are listed in Table 1.

**Table 1.** Antibodies used for flow cytometry of amniotic fluid. . This table provides specific information for each antibody used in multispectral profiling of the amniotic fluid.

| Marker | Fluorochrome | Clone | Source | Cat.# | RRID |
| --- | --- | --- | --- | --- | --- |
| CD11c | BUV563 | N418 | BD Horizon | 749040 | AB_2875243 |
| MHCII | BUV395 | 2G9 | BD Horizon | 569244 | AB_3684900 |
| CD19 | BV650 | 6D5 | BioLegend | 115541 | AB_11204087 |
| CD11b | BV570 | M1/70 | BioLegend | 101233 | AB_10896949 |
| IL-6 | PE-Cy7 | D7715A7 | BioLegend | 115814 | AB_2565575 |
| IL-31 | PE | W17037B | BioLegend | 160704 | AB_2876573 |
| NK-1.1 | PerCP/Cyanine5.5 | S17016D | BioLegend | 156525 | AB_2894655 |
| CD3 | SparkRed-718 | 17A2 | BioLegend | 100282 | AB_2924440 |
| Ly6c | PE-Dazzle-594 | HK1.4 | BioLegend | 128044 | AB_2566577 |
| CD45 | APC-Fire810 | 30-F11 | BioLegend | 103173 | AB_2860599 |
| CD335<br>(NKp46) | APC | 29A1.4 | BioLegend | 137608 | AB_10612758 |

### Amniotic fluid cytokine evaluation

Cytokines and chemokines were quantified from 20–30 µL of embryo-specific amniotic fluid using a 44-plex bead-based immunoassay (Luminex® xMAP®; MILLIPLEX® Mouse Cytokine/Chemokine 44-Plex, MCYTOMAG-70K), performed by Eve Technologies (Calgary, AB, Canada) on a Luminex® 200™ system. The panel measured inflammatory cytokines (e.g., TNF-α, IL-1α/β, IL-6, IL-10, IFN-γ), T-cell–associated cytokines (e.g., IL-2, IL-4, IL-5, IL-7, IL-9, IL-12p40/p70, IL-13, IL-15, IL-17), growth factors (e.g., G-CSF, GM-CSF, M-CSF, VEGF-A), and chemokines (e.g., CCL2/MCP-1, CCL3/MIP-1α, CCL4/MIP-1β, CCL5/RANTES, CCL11/Eotaxin, CXCL1/KC, CXCL2/MIP-2, CXCL9/MIG, CXCL10/IP-10, LIX). Concentrations were exported as CSV and processed in Python, non-numeric outputs were treated as missing and imputed using the cytokine-specific median. To harmonize analyte dynamic ranges, each cytokine was transformed using a robust Z-score; values were clipped to ±7.5 for visualization only. Heatmaps were generated per embryo (no litter averaging) and exported as vector PDFs.

### IL-6 blockade during MIA

Pregnant dams were assigned to saline, poly(I:C), or poly(I:C) + IL-6 blockade groups. For IL-6 blockade, dams received anti–mouse IL-6 (clone MP5-20F3, Cat# BE0046, RRID AB_1107709) at 8.3 mg/kg (166 µg for a 20 g dam; i.p.) 1 hour prior to poly(I:C) administration. Maternal serum and amniotic fluid were collected for IL-6 quantification as described above. Embryos were assessed for developmental abnormalities using standardized craniofacial and axial morphological metrics. Group differences were analyzed by one-way ANOVA with post hoc multiple-comparison testing, and representative images were acquired under identical settings.

## Statistical analyses

To ensure rigor, investigators were blinded to saline or poly(I:C) condition for all downstream analyses. Dams whose blood did not exhibit a significant increase in IL-6 3 hours after poly(I:C) injection were excluded from the study (this only affected about 2% of litters). In the majority of cases, experiments were performed between three and six times.

For all analyses, sample sizes were chosen based on previously generated data and/or power analyses. Acquired data was first tested for normality and log-normality before choosing a parametric or non-parametric statistical test. When the data were found to be normally distributed, parametric t-tests, one-way ANOVAs, or repeated measures two-way ANOVAs were used. If data were found to be non-Gaussian and non-logarithmic, a Mann-Whitney test was performed.

Statistical analyses were performed in Excel and Prism 9.0 (GraphPad Software). Figures were created using Python and Graphpad Prism and formatted using Adobe Illustrator (2024). The models were generated in biorender.com. Data are presented as mean ± SEM unless otherwise indicated.

## ACKNOWLEDGEMENTS

We thank Dr. Michael Wigler for insightful input at an early stage in the project. We also thank the following individuals for critical input and feedback on the manuscript: Dr. Corina Amor Vegas (CSHL), Dr. Jeremy Borniger (CSHL), Dr. Patrick Mitchell (University of Washington), Dr. Marty Yang (University of California in Los Angeles), Dr. Austin Ferro (CSHL), and Dr. Dominic Vita (CSHL). We also thank Life Science Editors for assistance with editing the final draft. We acknowledge the CSHL Cancer Center Microscopy Shared Resources for services and technical expertise (NCI 2P30CA045508). We also acknowledge the flow cytometry core facility and the histology core at CSHL. Lucas Cheadle is a Howard Hughes Medical Institute Freeman Hrabowski Scholar.

## Author contributions

Conceptualization: LC, ISM, BTK

Methodology: LC, ISM, BTK

Formal Analysis: ISM, BTK, BK, PH, DD

Investigation: ISM, BK, PH, QL, DD, VB, JP, BTK

Resources: VB, ISM, JP, LC

Validation: VB, ISM, BK, JP, LC

Data curation: ISM, BK, QL

Software: BK, QL

Writing – original draft: LC, ISM, BK

Writing – review & editing: LC, BTK, ISM, BK

Visualization: LC, ISM, BK, PH, QL

Supervision: LC, BTK

Funding acquisition: LC

Project administration: LC, BTK, JP

## Data, code, and materials availability

All data needed to evaluate and reproduce the results in this paper are present in the paper and/or the Supplementary Materials. This study did not generate new materials. Source code utilized in the study has been deposited to the Zenodo repository and can be found at https://zenodo.org/records/21709927?token=eyJhbGciOiJIUzUxMiJ9.eyJpZCI6IjJhYmRjYjY 2LTU5OTktNDI2Mi04MGZlLWEzMzZiZjBmODgwOSIsImRhdGEiOnt9LCJyYW5kb20iOi I2ZTQ4MDg2MWQzNjZjNDRiNDU3ZjQ5YzU0N2ZhZjE1NCJ9.BIa9MUnweDWSY2Coz kXv1CRmfaEExHydhujGRet0zAEopuXAq8XJMWvGka-HLLQ4mpFPSYc6CZyH2GZB5A509g. All snRNAseq data have been submitted to the Gene Expression Omnibus repository under accession number GSE315060.

