## Supplementary Materials for "Maternal-fetal immune conflict contributes to male-specific impairments in a mouse model of neurodevelopmental disorders"

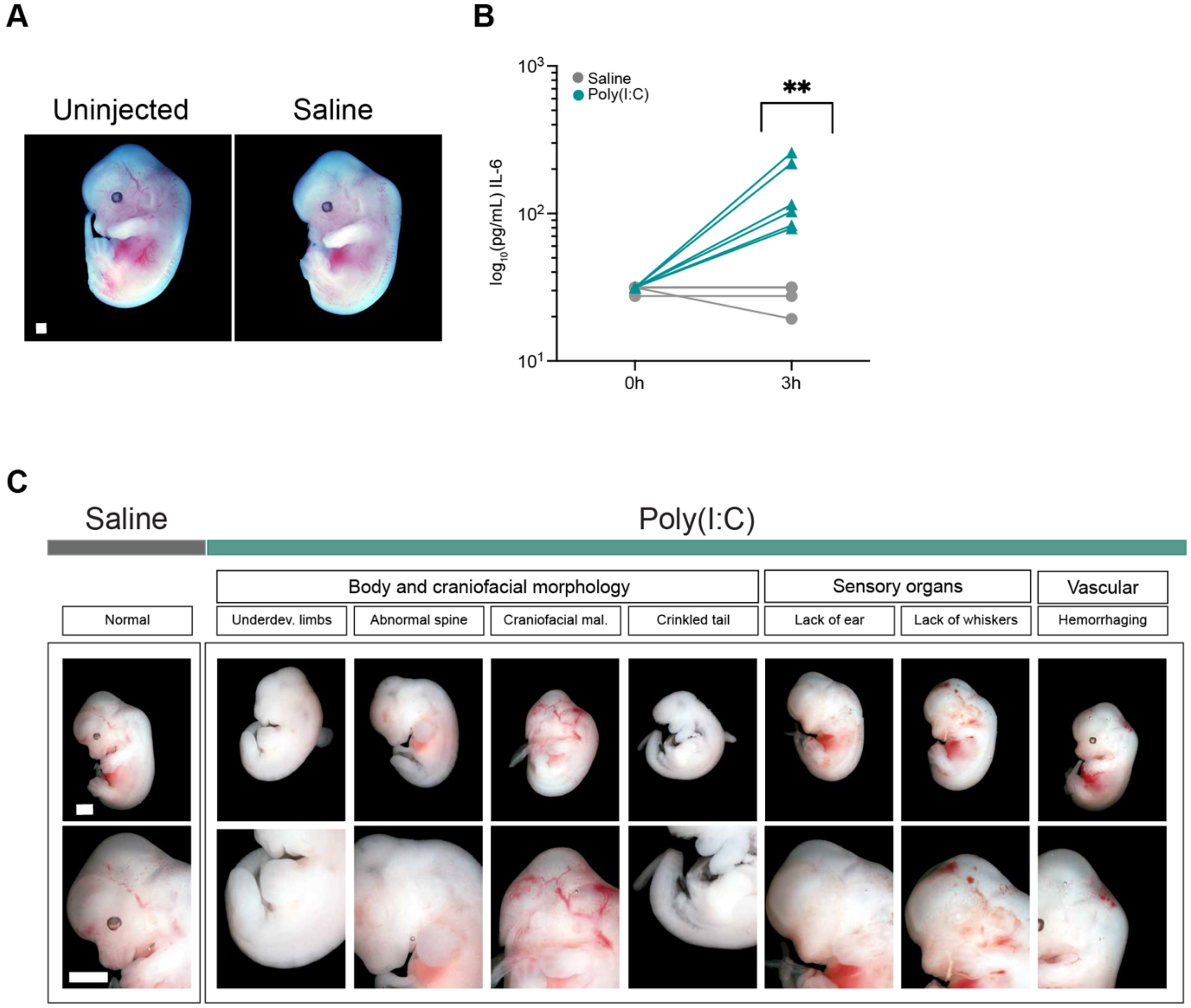


**Fig. S1. Characterization of MIA phenotypes in BALB/c mice**. (A) Example images of embryos at E13.5 from an uninjected dam and a dam injected with saline at E12.5. Scale bar, 0.9 mm. (B) Levels of IL-6 protein detected in the serum of BALB/c dams exposed to poly(I:C) (teal) or saline (gray) as measured by ELISA. Unpaired t test, $**p < 0.01$; n = 6 dams poly(I:C) and 4 dams saline. (C) Example images of embryos harvested from BALB/c dams at E13.5, 24 hours after saline or poly(I:C) exposure. Scale bars, 2 mm.

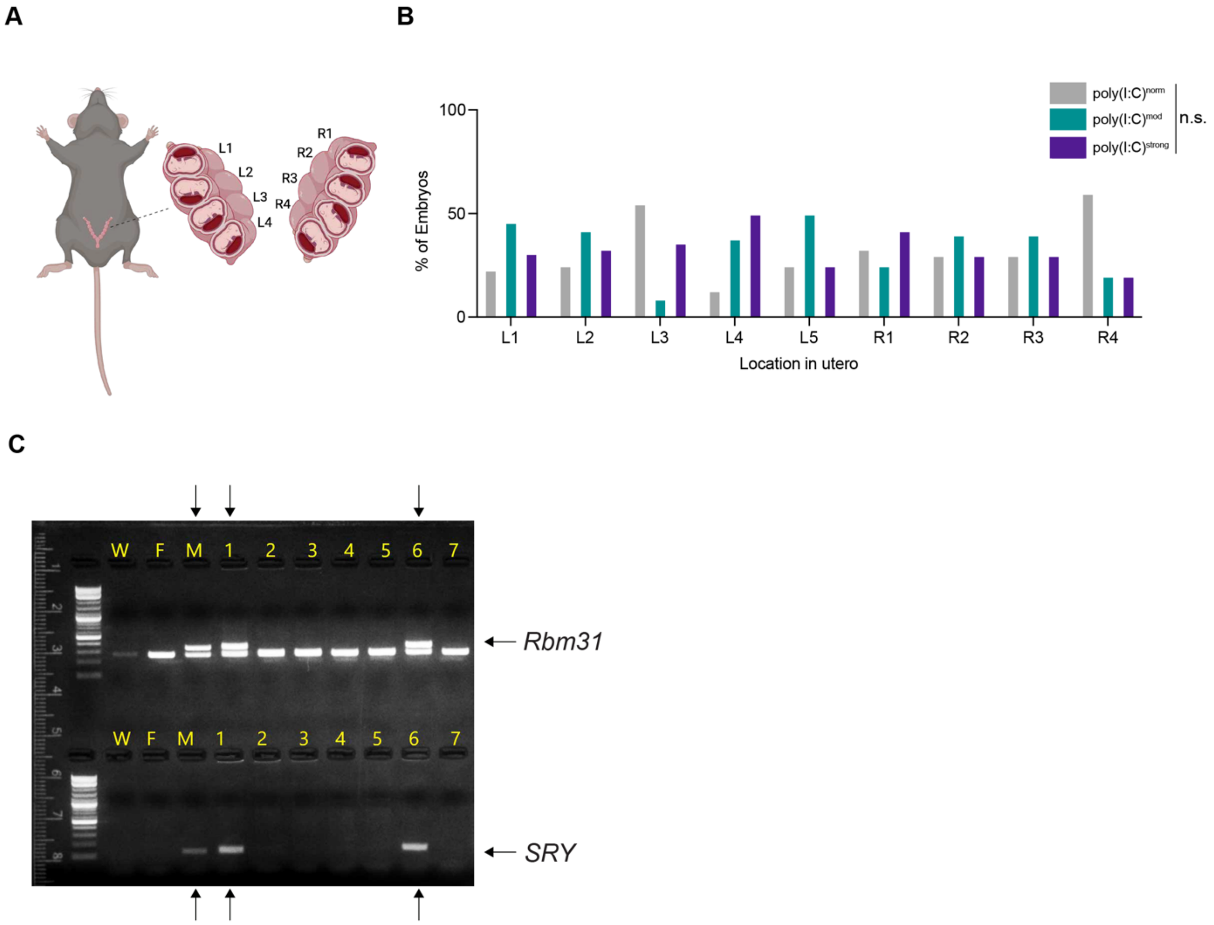


**Figure S2. Effects of intrauterine position and demonstration of sex determination**. (A) Schematic showing the organization of embryos within a pregnant dam. L, left uterine horn; R, right horn. Created in BioRender. Cheadle, L. (2026) https://BioRender.com/6pahnyu. (B) Percentage of poly(I:C)$^{norm}$, poly(I:C)$^{mod}$, and poly(I:C)$^{strong}$ embryos observed at each intrauterine position. Two-Way ANOVA with Tukey's post test: **p < 0.01; ***p < 0.001; and ****p < 0.0001; n =116 embryos from 15 distinct litters. (C) Example agarose gel demonstrating results of PCR genotyping to identify embryonic sex. *Rbm31*, upper band denotes the male copy. SRY, band reveals presence of the Y chromosome. Arrows, males.

A

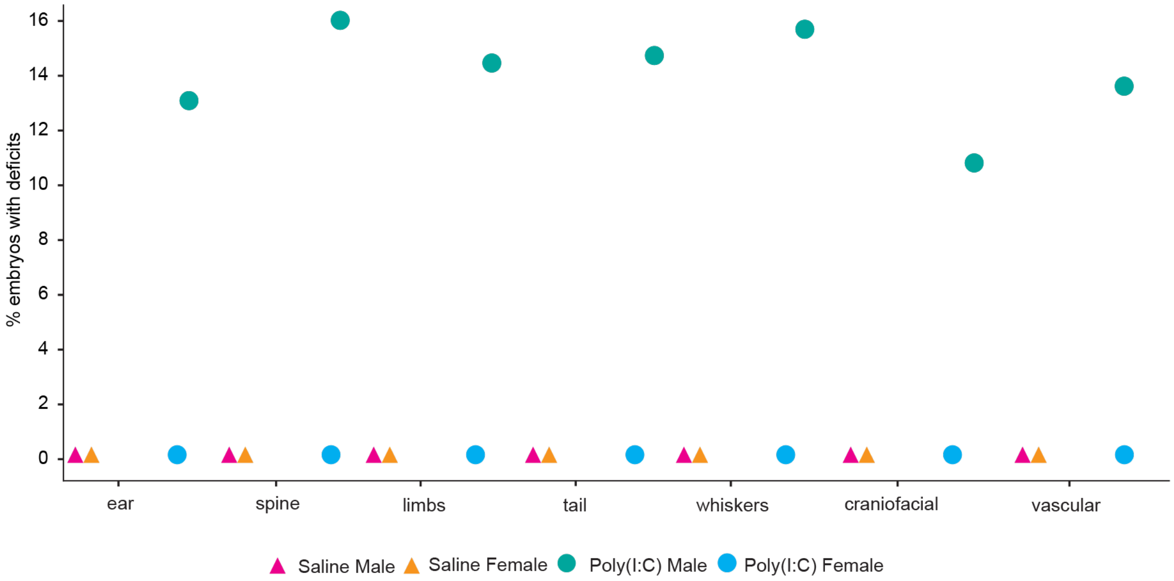


**Figure S3. MIA elicits developmental deficits in a subset of BALB/c male embryos**. Graph illustrating the percentage of embryos exhibiting deficits (x axis) across both sexes and poly(I:C) or saline treatment groups. Data aggregated from n = 43 embryos poly(I:C) and 36 embryos saline.

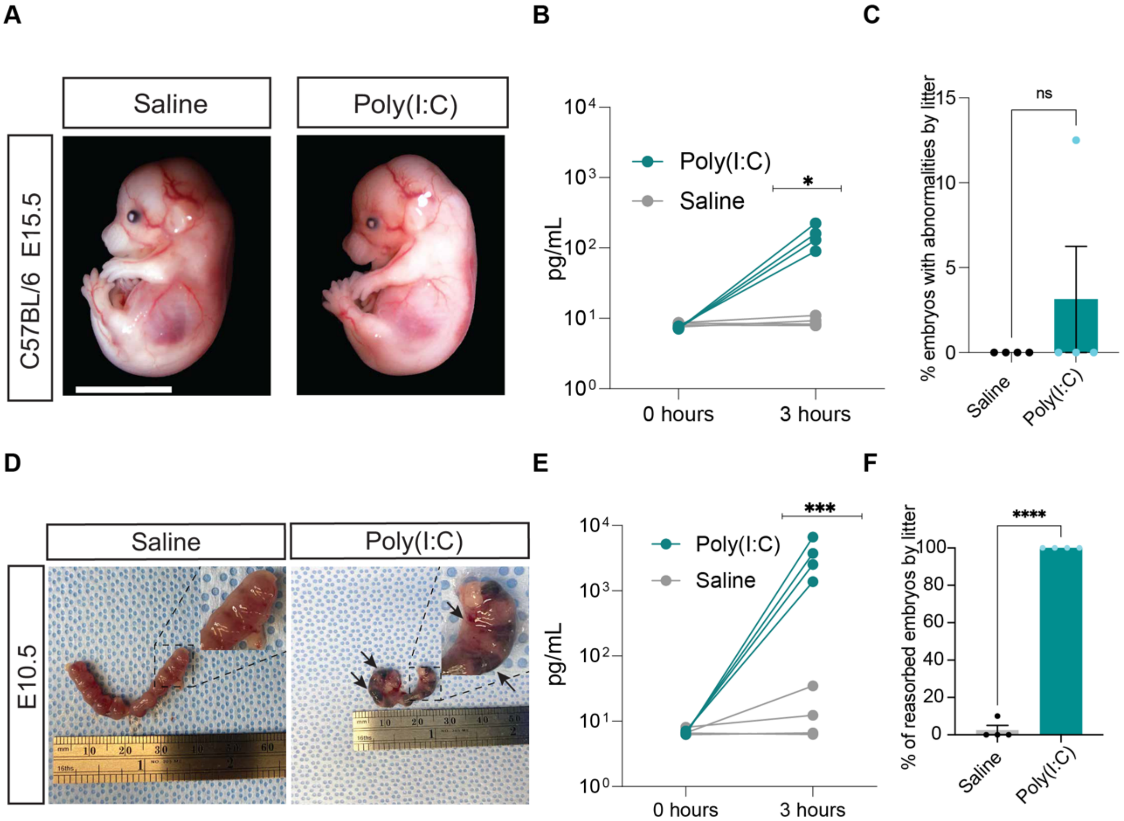


**Figure S4. E12.5 – E13.5 is a sensitive period of male susceptibility to MIA**. (A) Images of embryos harvested at E15.5 following saline or poly(I:C) injection at E14.5. Scale bar,10 mm. (B) Validation of IL-6 protein induction (ELISA) in the serum of dams injected with saline or poly(I:C) at E14.5. Unpaired t test, ***p < 0.001; n = 4 dams per condition. (C) Percentage of each litter exhibiting developmental abnormalities. Out of 22 embryos total, only one exhibited deficits following poly(I:C) injection at E14.5. Unpaired t test, n = 4 litters per condition. (D) Images of the conceptus within the uterine horns at E10.5 from dams injected with either saline or poly(I:C) at E9.5. All embryos from the poly(I:C)-treated group have hemorrhagic appearance indicating uterine demise and reabsorption. Scale bar, included in image. (E) Validation of IL-6 protein induction in dams injected at E9.5. with poly(I:C) (teal) or saline (gray), as measured by ELISA. Unpaired t test, *p < 0.05; n = 4 dams per condition. (F) Percentages of reabsorbed embryos by litter. Unpaired t test, ****p < 0.0001; n = 4 litters per condition.

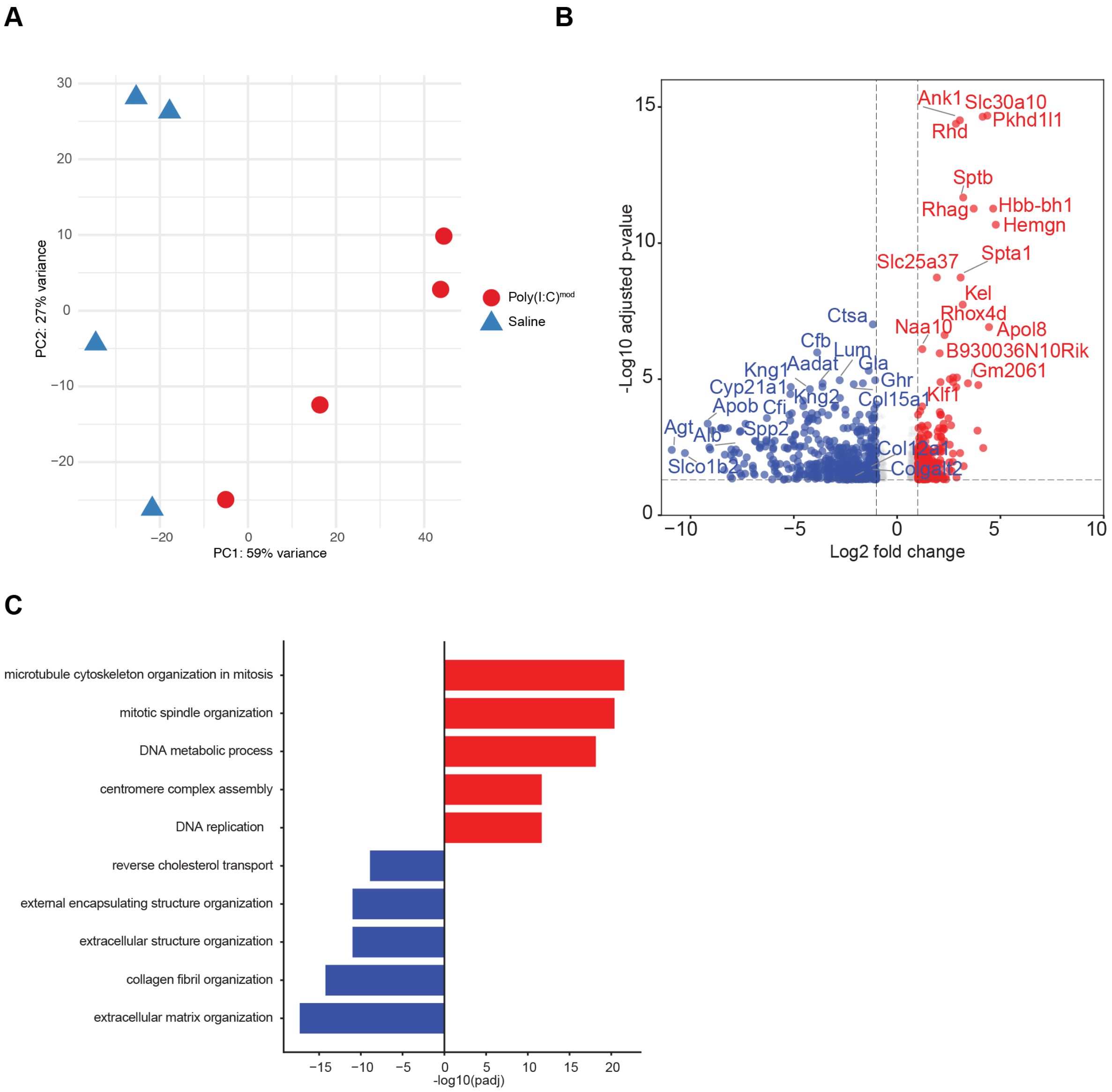


**Figure S5. MIA dampens extracellular matrix gene expression and enhances vascular-associated transcription in poly(I:C)$^{mod}$ placentas**. (A) Principal Component Analysis (PCA) reflecting separation between saline and poly(I:C) samples based on placental bulk RNA-sequencing; n = 4 placentas per condition. (B) Volcano plot illustrating transcripts that were upregulated (red) or downregulated (blue) in poly(I:C)$^{mod}$ versus saline placentas, as measured by RNA seq. (C) Gene ontology analysis demonstrating enriched functional categories of differentially expressed genes between the two conditions. Red, GO categories enriched among genes upregulated in poly(I:C)$^{mod}$ placentas versus saline; blue, GO categories enriched among genes downregulated in poly(I:C)$^{mod}$ placentas versus saline.

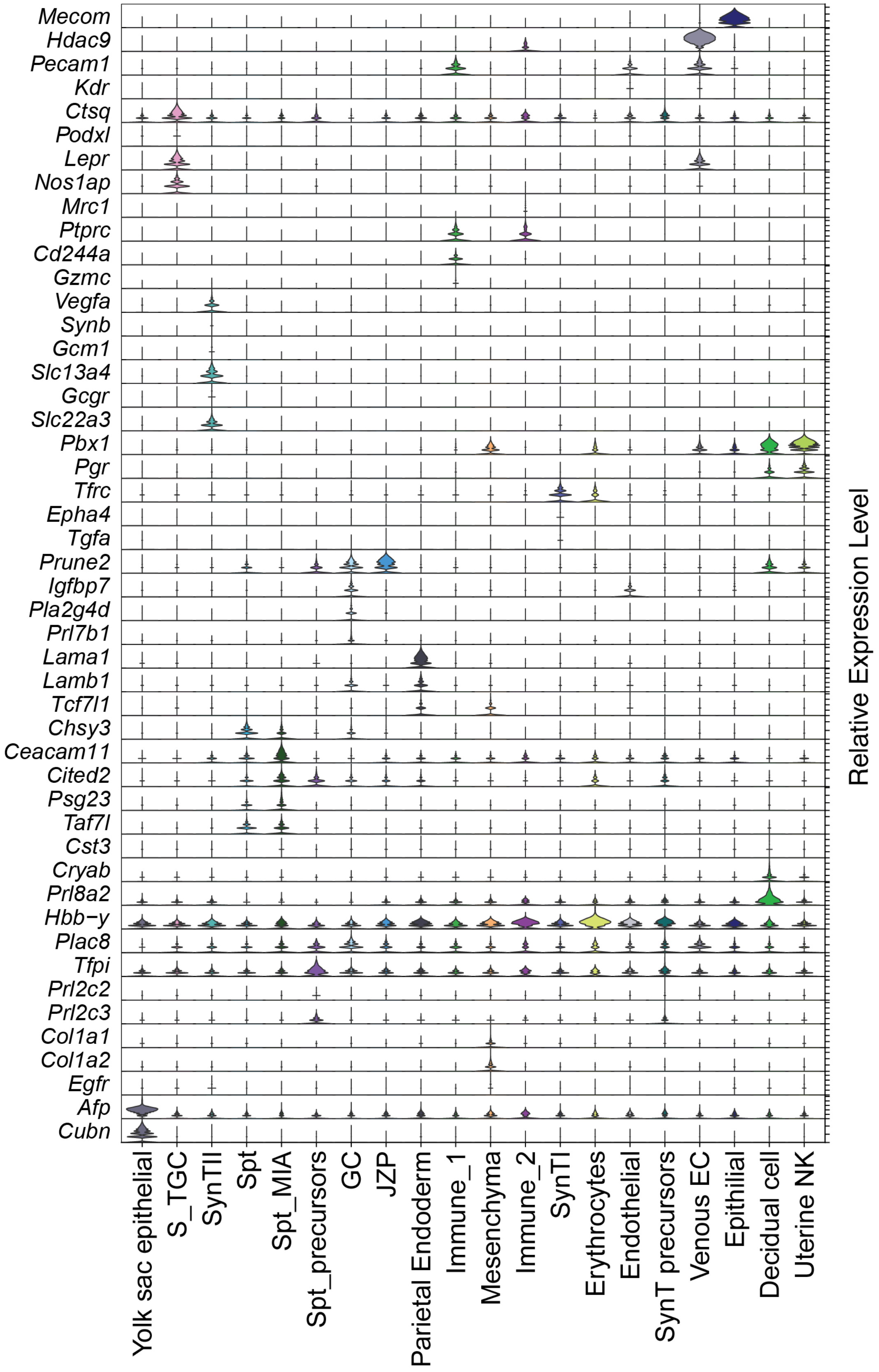
Mecom
Hdac9
Pecam1
Kdr
Ctsq
Podxl
Lepr
Nos1ap
Mrc1
Ptprc
Cd244a
Gzmc
Vegfa
Synb
Gcm1
Slc13a4
Gcgr
Slc22a3
Pbx1
Pgr
Tfrc
Epha4
Tgfa
Prune2
Igfbp7
Pla2g4d
Prl7b1
Lama1
Lamb1
Tcf7l1
Chsy3
Ceacam11
Cited2
Psg23
Taf7l
Cst3
Cryab
Prl8a2
Hbb−y
Plac8
Tfpi
Prl2c2
Prl2c3
Col1a1
Col1a2
Egfr
Afp
Cubn
Relative Expression Level
Yolk sac epithelial
S_TGC
SynTII
Spt
Spt_MIA
Spt_precursors
GC
JZP
Parietal Endoderm
Immune_1
Mesenchyma
Immune_2
SynTI
Erythrocytes
Endothelial
SynT precursors
Venous EC
Epithilial
Decidual cell
Uterine NK

**Figure S6. Cell type annotations of placental cells defined by single-nucleus RNA sequencing**. Violin plot demonstrating expression of enriched genes across cell types, serving as cell type markers for the snRNAseq dataset.

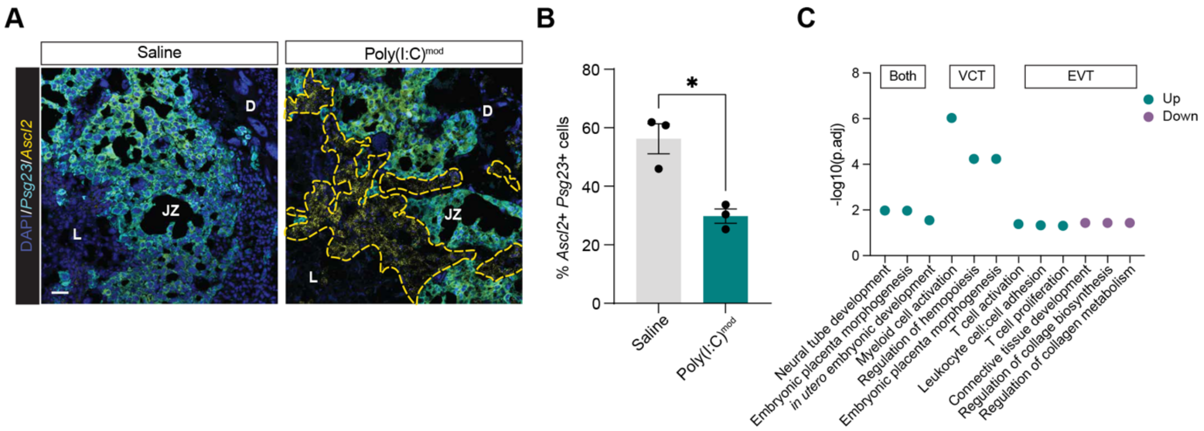


**Figure S7. Decreased overall *Psg23* expression in spongiotrophoblasts and comparison of our data with a human dataset**. (A) Confocal images of placental sections from saline-treated and poly(I:C)$^{mod}$ male embryos subjected to fluorescence *in situ* hybridization for the spongiotrophoblast (SpT) marker *Ascl2* (yellow) and *Psg23* (cyan). Yellow outline, SpTs not expressing *Psg23*. D, decidua; JZ, junctional zone; L, labyrinth. Scale bar, 50 μm. (B) Quantification of the percentage of cells double-positive for *Ascl2* and *Psg23*. Unpaired t-test, $*p < 0.05$, n=3 placentas per condition. (C) Gene ontology (GO) analysis of genes up- or downregulated in SpTs following MIA that are also dysregulated in villous cytotrophoblasts (VCTs) or extravillous trophoblasts (EVTs) in human placentas exposed to COVID-19 during late pregnancy. GO terms shared among upregulated genes, teal; GO terms shared among downregulated genes, purple.

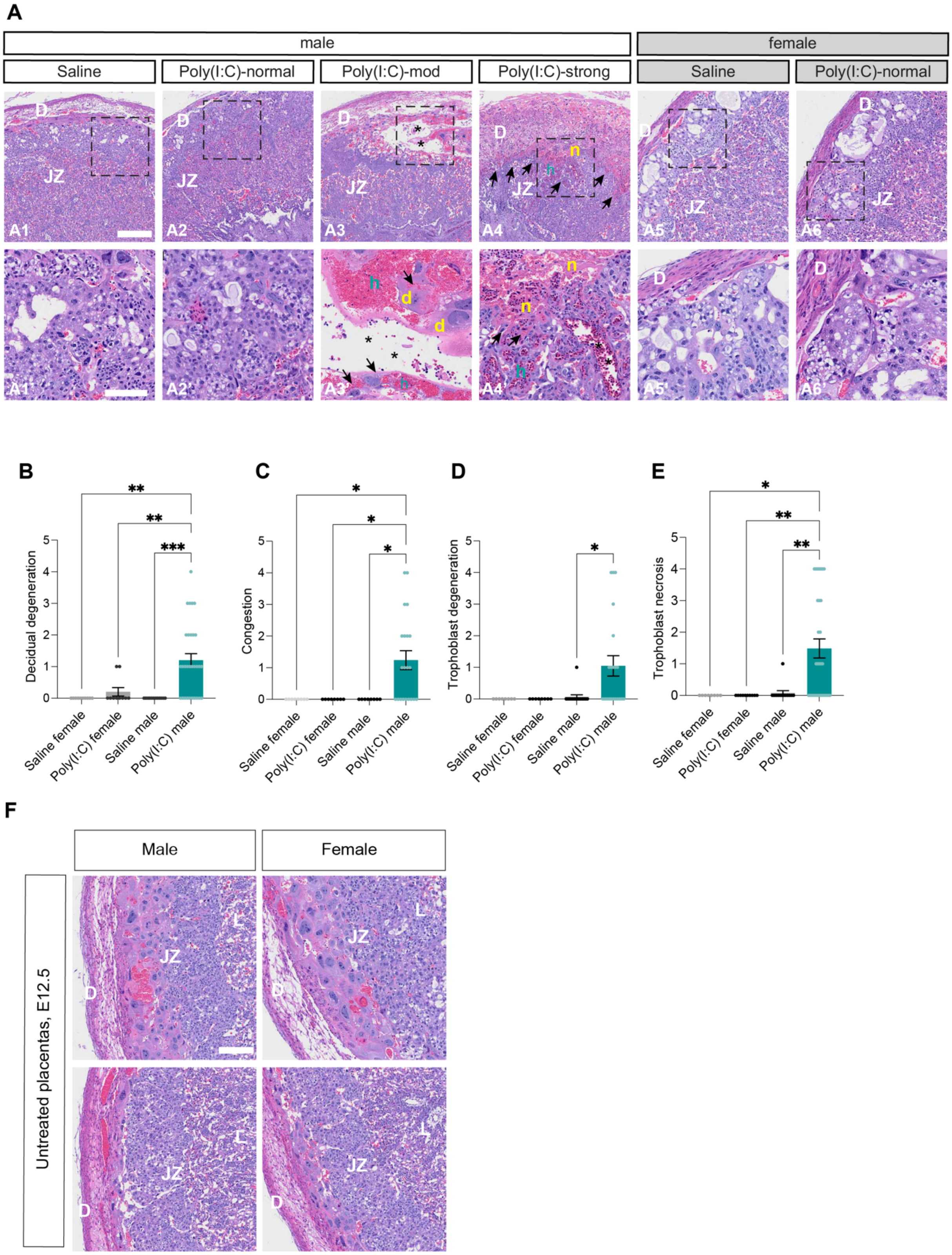

A
male
female
Saline
Poly(I:C)-normal
Poly(I:C)-mod
Poly(I:C)-strong
Saline
Poly(I:C)-normal
B
Decidual degeneration
C
Congestion
D
Trophoblast degeneration
E
Trophoblast necrosis
Saline female
Poly(I:C) female
Saline male
Poly(I:C) male
F
Male
Female
Untreated placentas, E12.5

**Figure S8. Breakdown in placental integrity is restricted to poly(I:C)$^{mod}$ and poly(I:C)$^{strong}$ males**. (A) Example images of placental sections subjected to H&E staining across sexes, conditions, and phenotypes. D, decidua; JZ, junctional zone; d, degeneration; h, hemorrhagic congestion; n, necrosis. Scale bar, 500 µm. Inset scale bar, 100 µm. (B)-(E) Quantification of different degenerative signatures observed by H&E staining. Data represent severity scores determined by a pathologist blinded to condition. Decidual degeneration (B), congestion (C), trophoblast degeneration (D), and trophoblast necrosis (E). Two-way ANOVAs with Tukey's post test; *$p < 0.05$, **$p < 0.01$; ***$p < 0.001$; n = 10 placentas per condition. (F) Example images of E12.5 placental sections from male and female embryos subjected to H&E staining. Scale bar, 500 µm.

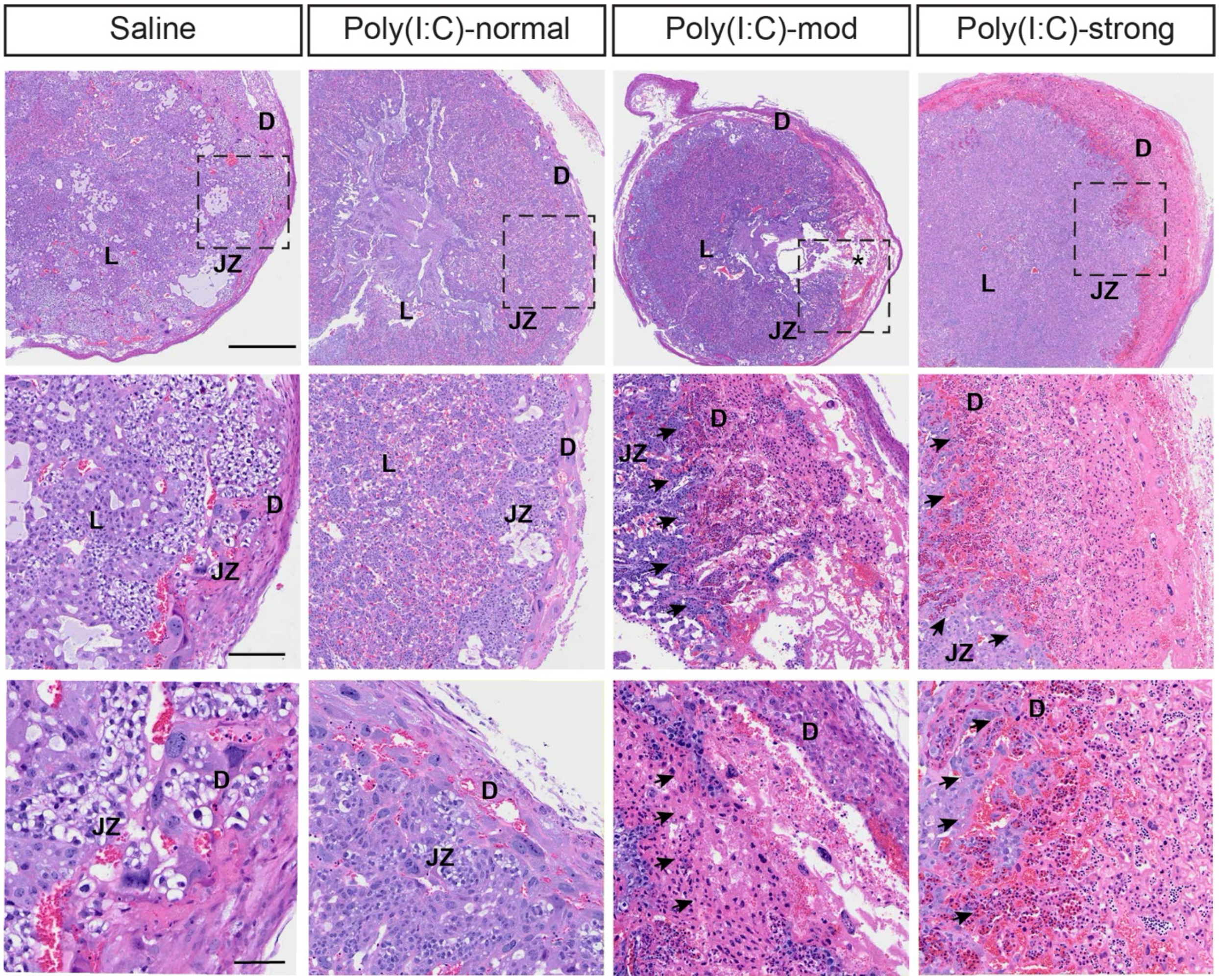


**Figure S9. Examples of decidual pathology in the placentas of male mice across conditions and phenotypes.** Low-, medium-, and high-magnification images of decidual pathology at the maternal–fetal interface following MIA based upon H&E staining of placental sections. D, decidua; JZ, junctional zone; and L, labyrinth. Note the presence of vascular congestion and hemorrhagic areas, expansion of interstitial spaces consistent with edema, loss of normal tissue organization, eosinophilic regions with reduced cellularity suggestive of necrotic degeneration, and inflammatory cell accumulation (arrowheads), all signs of decidual degeneration observed only in poly(I:C)$^{mod}$ and poly(I:C)$^{strong}$ male placentas. Dashed boxes indicate regions shown at higher magnification in the panels below. Scale bar, 500 µm (top row). Inset scale bars, 100 µm (middle row) and 50 µm (bottom row).

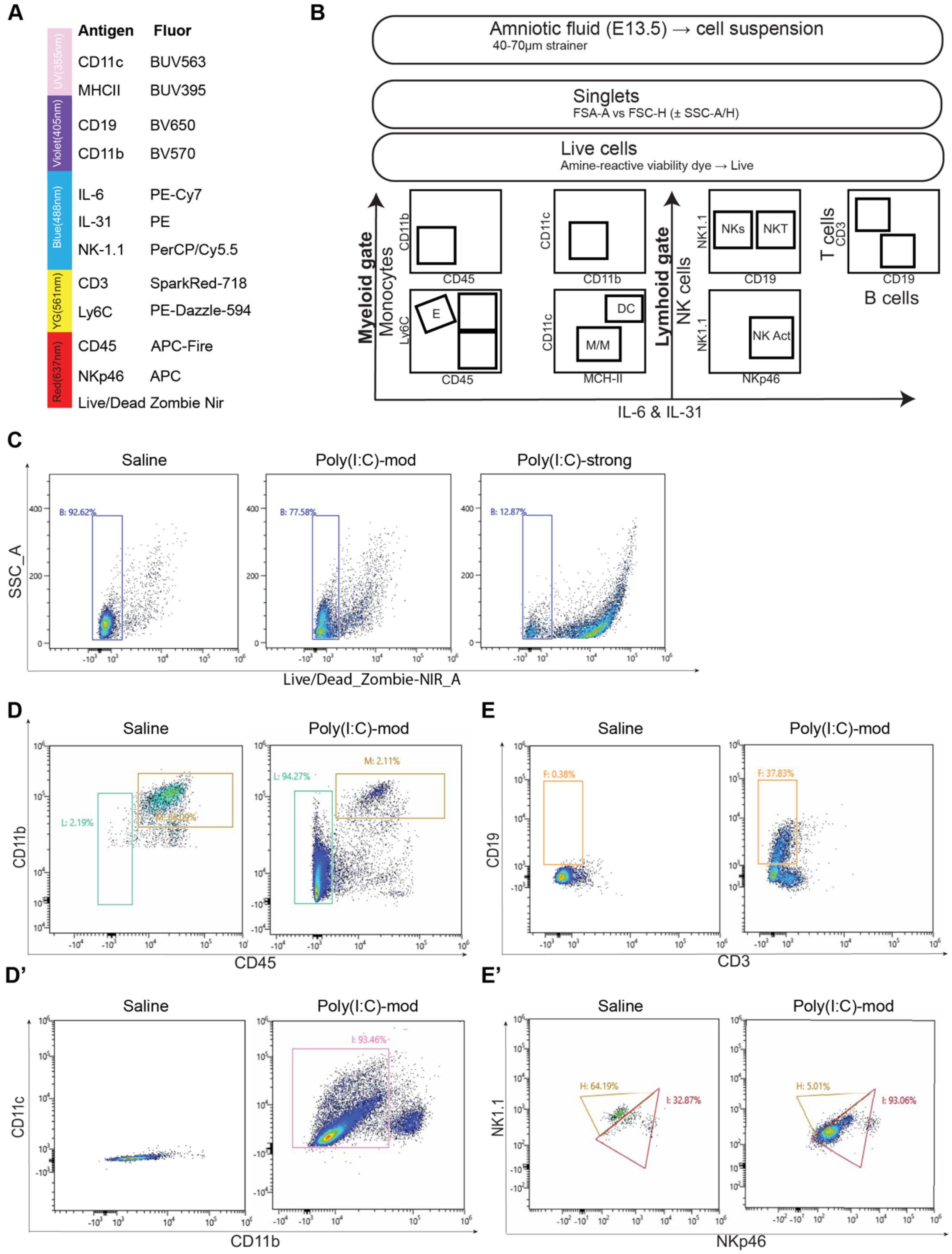
A
Antigen Fluor
CD11c BUV563
MHCII BUV395
CD19 BV650
CD11b BV570
IL-6 PE-Cy7
IL-31 PE
NK-1.1 PerCP/Cy5.5
CD3 SparkRed-718
Ly6C PE-Dazzle-594
CD45 APC-Fire
NKp46 APC
Live/Dead Zombie Nir
UV(355nm)
Violet(405nm)
Blue(488nm)
YG(561nm)
Red(637nm)
B
Amniotic fluid (E13.5) → cell suspension
40-70µm strainer
Singlets
FSA-A vs FSC-H (± SSC-A/H)
Live cells
Amine-reactive viability dye → Live
Myeloid gate
Monocytes
Lymhoid gate
NK cells
T cells
B cells
NKs
NKT
DC
M/M
NK Act
IL-6 & IL-31
C
Saline
Poly(I:C)-mod
Poly(I:C)-strong
B: 92.62%
B: 77.58%
B: 12.87%
SSC_A
Live/Dead_Zombie-NIR_A
D
E
L: 2.19%
L: 94.27%
M: 2.11%
F: 0.38%
F: 37.83%
CD11b
CD45
CD19
CD3
D'
E'
I: 93.46%
H: 64.19%
I: 32.87%
H: 5.01%
I: 93.06%
CD11c
NK1.1
NKp46

**Figure S10. Multiplexed flow cytometric strategy for quantifying immune cells within the amniotic fluid.** (A) Antibody/fluorochrome panel. (B) Schematic of the gating strategy. E13.5 amniotic fluid was filtered to obtain a single-cell suspension, then gated for singlets and live cells, followed by separation of myeloid and lymphoid compartments. (C–E′) Representative plots illustrating sequential gating, including live-cell selection (C), CD45 vs. CD11b leukocyte/myeloid identification (D), CD11b vs. CD11c subdivision (D′), CD3 vs CD19 T/B cell gating (E), and NK1.1 vs NKp46 NK/NKT-related subsets (E′). Numbers indicate the percentage of events within each gate for the representative samples shown.

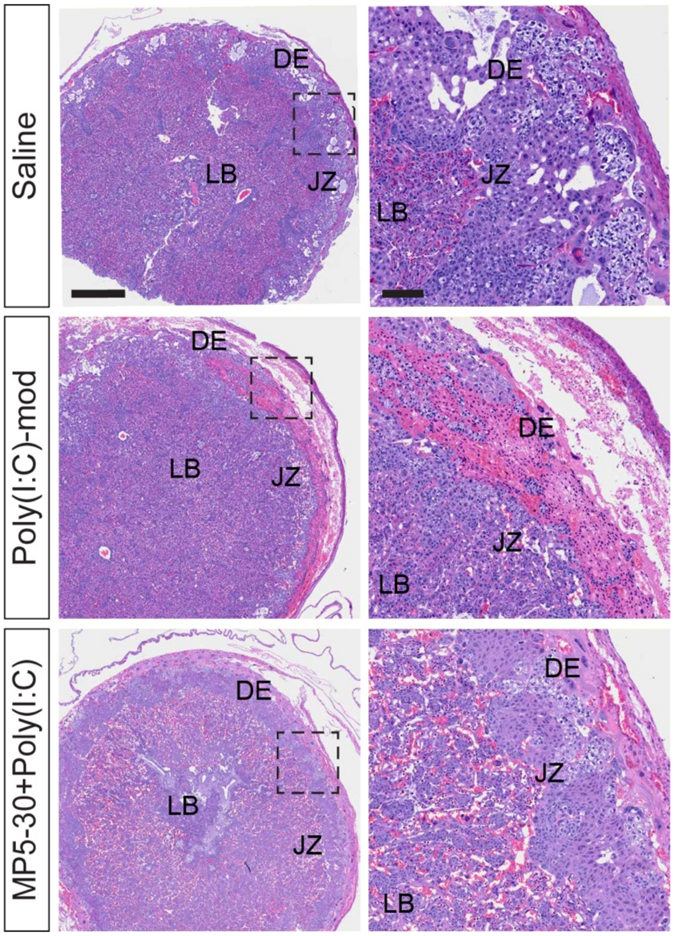


**Figure S11. Neutralization of IL-6 maintains placental architecture following MIA**. H&E-stained placental sections harvested from embryos exposed to poly(I:C) with and without neutralization of IL-6. DE, decidua; JZ, junctional zone; and LB, labyrinth. Dashed boxes indicate inset magnifications on the right. Scale bar, 500 µm. Inset scale bar, 100 µm.

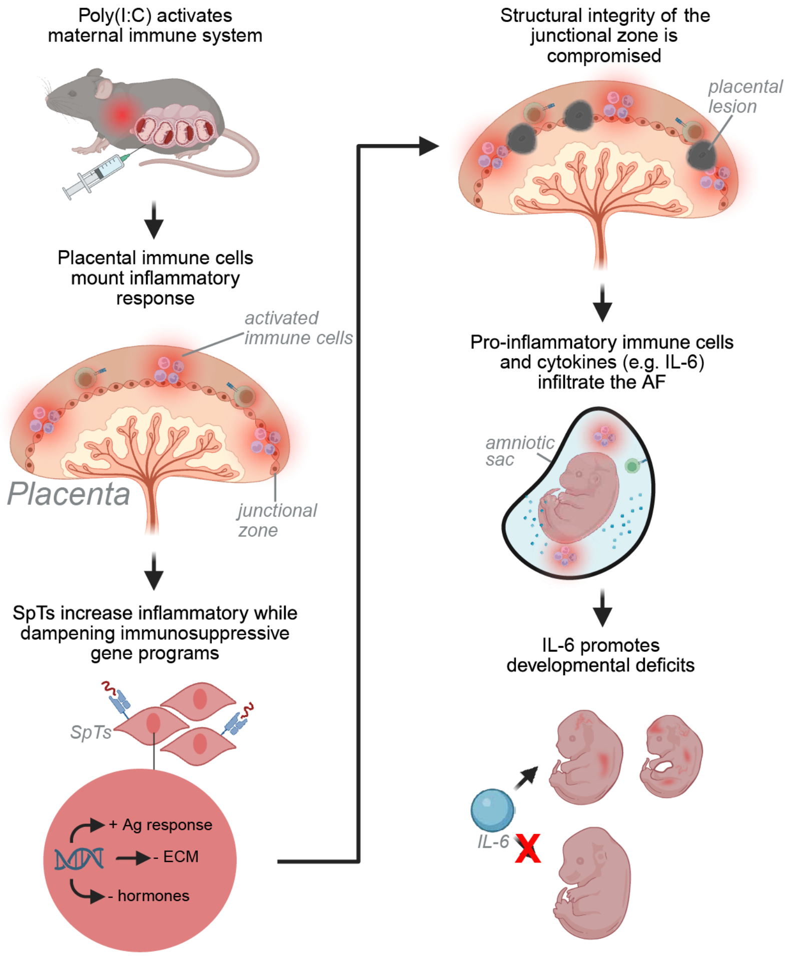


**Figure S12. Impact of MIA on a subset of male embryos in the MIA$_{\text{poly(I:C)}}$ mouse model.** Poly(I:C) induces an immune response in the dam leading to inflammation within the placentas of a subset of male embryos. Fetally derived spongiotrophoblasts which lie at the border of the maternal and fetal compartments of the placenta exhibited an increase in inflammatory gene expression, concurrent with a decrease in extracellular matrix and hormone-associated transcripts. As a result, the structural integrity of the placenta is compromised, contributing to the infiltration of immune cells and cytokines into the amniotic fluid. One of these cytokines, IL-6, is required for the emergence of placental and fetal deficits in male offspring. Created in BioRender. Cheadle, L. (2026) https://BioRender.com/zthdjda.

| | 2 Way ANOVA: | | | Tukey's post test: | |
|---|---|---|---|---|---|
| Cytokine | Interaction | sex | treatment | saline male vs. poly(I:C) male | saline female vs. poly(I:C) female |
| TNFa | *** | *** | *** | **** | ns |
| Gm-CSF | **** | *** | ** | **** | ns |
| MIP-2 | ** | ** | ** | **** | ns |
| IL-1b | ** | ns | ns | ** | ns |
| IL-6 | *** | *** | *** | **** | ns |
| IL-17 | * | ns | * | * | ns |
| IL-2 | *** | ns | ns | *** | ns |
| IL-3 | * | **** | ns | ns | ns |
| IL-5 | * | * | * | ** | ns |
| IL-11 | * | * | ** | *** | ns |
| LIF | **** | *** | **** | **** | ns |
| KC | ns | ** | ** | ** | ns |
| MCP-1 | ns | ** | ** | * | ns |
| IP-10 | ** | *** | ** | **** | ns |

**Table S1. Statistics of cytokine abundance in the amniotic fluid of females and males exposed to saline or poly(I:C)**. Statistics describing the abundance of cytokines in amniotic fluid of males and females. All data were analyzed using a 2-Way ANOVA followed by Tukey's post test. *$p < 0.05$; **$p < 0.01$; ***$p < 0.001$; ****$p < 0.0001$; ns $p > 0.05$.
